# Cargo genes of Tn7-like transposons comprise an enormous diversity of defense systems, mobile genetic elements and antibiotic resistance genes

**DOI:** 10.1101/2021.08.23.457393

**Authors:** Sean Benler, Guilhem Faure, Han-Altae Tran, Sergey Shmakov, Feng Zheng, Eugene Koonin

## Abstract

Transposition is a major mechanism of horizontal gene mobility in prokaryotes. However, exploration of the genes mobilized by transposons (cargo) is hampered by the difficulty in delineating integrated transposons from their surrounding genetic context. Here, we present a computational approach that allowed us to identify the boundaries of 6,549 Tn7-like transposons at base pair resolution. We found that 96% of these transposons carry at least one cargo gene. Delineation of distinct communities in a gene-sharing network demonstrates how transposons function as a conduit of genes between phylogenetically distant hosts. Comparative analysis of the cargo genes reveals significant enrichment of mobile genetic elements (MGEs) nested within Tn7-like transposons, such as insertion sequences and toxin-antitoxin modules, genes involved in recombination and anti-MGE defense, and in antibiotic resistance. More unexpectedly, cargo also includes genes encoding central carbon metabolism enzymes. Twenty-two Tn7-like transposons carry both an anti-MGE defense system and antibiotic resistance genes, illustrating how bacteria can overcome these combined pressures upon acquisition of a single transposon. This work substantially expands the distribution of Tn7-like transposons, defines their evolutionary relationships and provides a large-scale functional classification of prokaryotic genes mobilized by transposition.

**Significance:** Transposons are major vehicles of horizontal gene transfer that, in addition to genes directly involved in transposition, carry cargo genes. However, characterization of these genes is hampered by the difficulty of identification of transposon boundaries. We developed a computational approach for detecting transposon ends and applied it to perform a comprehensive census of the cargo genes of Tn7-like transposons, a large class of bacterial mobile genetic elements (MGE), many of which employ a unique, CRISPR-mediated mechanism of site-specific transposition. The cargo genes encompass a striking diversity of MGE, defense and antibiotic resistance systems. Unexpectedly, we also identified cargo genes encoding metabolic enzymes. Thus, Tn7-like transposons mobilize a vast repertoire of genes that can have multiple effects on the host bacteria.

## Introduction

Horizontal gene transfer (HGT) between prokaryotic genomes is one of the principal forces shaping prokaryotic genome evolution (Koonin, 2015; Nelson-Sathi et al., 2015; Puigbò et al., 2014; Sela et al., 2019; Takeuchi et al., 2014; Treangen and Rocha, 2011). The major routes of HGT include transformation, transduction, conjugation and transposition. Although the contribution of each route to the total number of horizontally transferred genes in a given genome is an outstanding question (Halary et al., 2010; McInnes et al., 2020; Popa et al., 2017), examining the genes mobilized via each pathway offers the opportunity to identify trends universal to all routes. Recent large-scale analyses of horizontally transferred genes have either not considered the molecular pathway of transfer (Oliveira et al., 2017; Song et al., 2019; Yaffe and Relman, 2020) or focused on a particular class of mobile genetic elements (MGE), such as plasmids (Antipov et al., 2019; Redondo-Salvo et al., 2020), integrative and conjugative elements (Cury et al., 2017) or viruses (Benler et al., 2020; Breitbart et al., 2018; Roux et al., 2020). However, to the best of our knowledge, the diversity of genes mobilized via transposition has not been comprehensively characterized.

In prokaryotes, transposons that mobilize using a DDE/RNaseH family transposase (e.g., insertion sequences) are the most abundant class (Siguier et al., 2015). At a minimum, an autonomous DNA transposon encodes a transposase and conserved transposase binding sites at the 5’ and 3’ ends of the element, where the transposase executes strand cleavage and subsequent transfer of the element from the donor to the acceptor site (Craig, 2015). Some DDE/RNaseH transposon families also require genes involved in regulatory and target site-selection steps (Hickman and Dyda, 2015), such as the model transposon Tn7 from *Escherichia coli.* Detailed biochemical dissection of Tn7 identified five “core” genes that collectively execute transposon excision, target site-selection and integration (Peters and Craig, 2001; Waddell and Craig, 1988). The Tn7 transposase, TnsB, recognizes arrays of binding sites at the left and right ends of the element and mediates 3’ DNA strand cleavage (Arciszewska et al., 1989; Peters and Craig, 2001). A second transposase, TnsA, mediates cleavage of the 5’ end that results in the complete excision of the element from the donor site (Sarnovsky et al., 1996). Transposition is orchestrated by TnsC, an AAA-ATPase that interposes between the TnsAB proteins bound to the ends of the transposon and the target site-selecting protein (Bainton et al., 1993; Stellwagen and Craig, 1997). The target site is recognized either by TnsD, a site-specific DNA-binding protein, or TnsE, a DNA-binding protein that directs integration into replicating DNA (Bainton et al., 1993; Parks et al., 2009).

Recently, it has been shown that some Tn7-like transposons recognize their target sites via a CRISPR spacer-guided mechanism (Hsieh and Peters, 2021; Klompe et al., 2019; Petassi et al., 2020; Saito et al., 2021; Strecker et al., 2019). Such CRISPR associated transposons (CASTs) encode either a subtype I-F or subtype I-B Cascade complex, or an inactivated type V-K effector (Faure et al., 2019; Peters et al., 2017). Another distinguishing feature of CASTs is an “atypical” repeat and spacer delocalized from the CRISPR array, which form the guide RNAs directing the transposon to the chromosomal target site (Hsieh and Peters, 2021; Petassi et al., 2020; Saito et al., 2021; Strecker et al., 2019). The site selectivity and regulation of transposition by Tn7 and by the CASTs stand in stark contrast to the near random insertion by other DDE-family transposons (Siguier et al., 2015).

In addition to the core genes of Tn7 and CASTs, other “cargo” genes that are not involved in transposition are often present within the boundaries of transposons (Parks and Peters, 2007; Peters and Craig, 2001). Along with the core genes, the cargo genes are mobilized from a donor site to a recipient site, making a substantial contribution to HGT (Parks and Peters, 2009; Peters et al., 2014). Examination of the cargo carried by Tn7 and about 50 other Tn7-like transposons identified integrons with antibiotic resistance gene cassettes, heavy metal resistance genes, iron-sequestering siderophores, non-ribosomal peptide synthases, restriction-modification enzymes and many other genes of unknown function (Aprile et al., 2021; Hamidian and Hall, 2021; Parks and Peters, 2007, 2009). Therefore, a larger-scale analysis of the cargo carried by Tn7-like transposons has the potential to illuminate consistent trends in transposon-mediated HGT.

A challenge for the study of integrated MGEs, including Tn7-like transposons, is accurate delineation of the 5’ and 3’ boundaries of the element. Several bioinformatic tools delimit integrated MGEs by identifying a local enrichment of MGE-associated gene annotations (e.g., integrases) in a given locus (Antipov et al., 2019; Antipov et al., 2020; Cury et al., 2020; Liu et al., 2018; Nayfach et al., 2020; Saak et al., 2020). However, because the diversity of genes carried by transposons is unknown, this approach is circular and thus of limited utility. Other tools delimit MGE boundaries by aligning sequence reads from the query genome against a reference (Durrant et al., 2020; Jiang et al., 2019), but selecting a reference genome is non-trivial. Thus, to characterize Tn7-like transposons on a large scale, it is highly desirable to develop an approach that is agnostic to gene functions and does not require a closely related reference genome or pangenome.

Here, we present a comprehensive survey of Tn7-like transposons in prokaryotic genomes and whole-community metagenomes. The conserved sequences of the transposase binding sites and their unique architecture are shown to carry a signal that is sufficient to delineate the 5’ and 3’ boundaries of the transposons, unmasking the diverse repertoire of genes mobilized by these elements. The comprehensive dissection of Tn7-like transposons enabled by this approach provides insight into prokaryotic HGT by expanding the known phyletic range of Tn7-like transposons, assessing the preferred routes and phylogenetic barriers to transposition, and characterizing the diversity of the mobilized genes.

## Results

### Tn7-like transposons are present across diverse bacterial phyla

To investigate the distribution of Tn7-like transposons in prokaryotic genomes, HMMs for the core genes of the *E. coli* Tn7 (*tnsABCDE)* and known Tn7-like transposons were used to identify homologs in the database of prokaryotic genomes. Loci were considered candidate Tn7- like transposons if they encompassed adjacent ORFs with significant sequence similarity to the core transposase TnsB and at least one other Tn7 HMM. The TnsB transposase sequences were employed for phylogenetic reconstruction, given that TnsB is required for transposition whereas the other subunit of the heteromeric transposase, TnsA, is dispensable (May and Craig, 1996). Using an iterative procedure to construct the alignments (Wolf et al., 2018), a single TnsB protein sequence alignment was employed to construct a comprehensive phylogenetic tree of Tn7-like transposons (Fig 1A).

**Figure 1.**
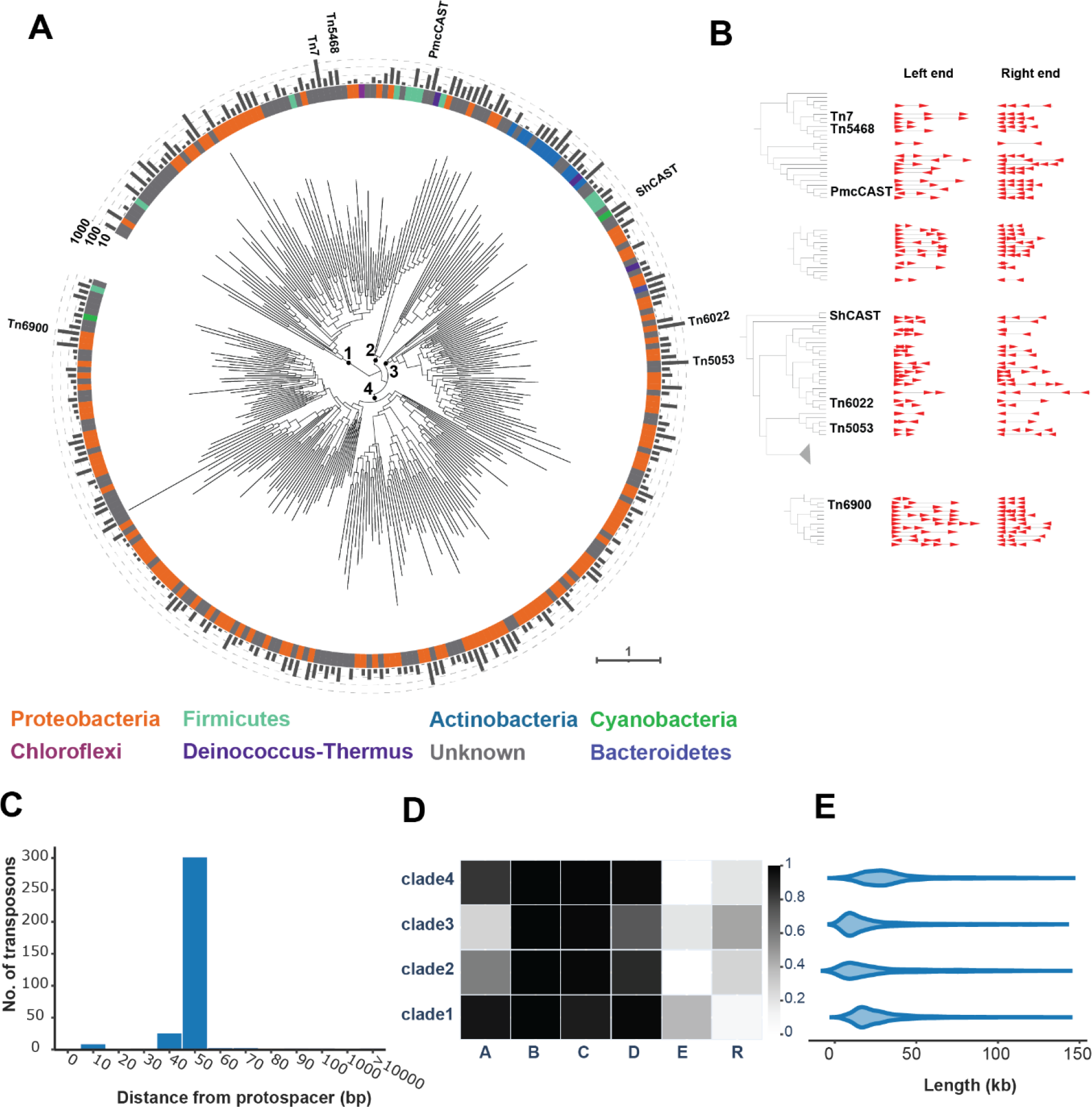
Tn7-like transposons are widespread mobile genetic elements across the diversity of bacteria. Phylogenetic tree of the DDE-family TnsB transposase from Tn7 and related transposons (n = 299 leaves). Deep branches are marked for clarity and the number of transposases represented by each leaf are plotted on the outer ring. The taxonomic phylum for each leaf is plotted on the inner ring and assigned using the consensus of ≥80% of the transposase sequences; otherwise, the leaf phylum is marked as “unknown” (**A**). The predicted boundaries from selected transposons are displayed for a TnsB subtree, highlighting related transposases possess similar TnsB binding site architectures. Each binding site is depicted as an arrow, where sites downstream of the transposase were arbitrarily oriented to the positive strand (left end, arrows pointing left-to-right) and sites upstream of the transposase are on the negative strand (right end, arrows pointing right-to-left) (**B**). Histogram of distances from the predicted boundary of CASTs to the protospacer-targeted attachment site (**C**). Heatmap of the presence/absence of Tn7 core proteins (**D**). Length distribution of intact, dereplicated transposons (n = 6,549) **(E).**

The tree partitioned into four well-supported clades of transposons (bootstrap support > 80) and the leaves from each clade were assigned a taxonomic host phylum using an 80% consensus rule. The first clade includes Tn7, an experimentally-characterized subtype I-B CAST (Saito et al., 2021) and other related transposons in the phyla Proteobacteria, Firmicutes and Chloroflexi. The second clade is dominated by Actinobacteria transposons, followed by transposons in the Proteobacteria, Firmicutes and Deinococcus-Thermus phyla, none of which have been experimentally characterized. The third clade includes the Proteobacteria transposons Tn5053 (Kholodii et al., 1995) and Tn6022 (Hamidian and Hall, 2011), Cyanobacterial subtype V-K CASTs (Strecker et al., 2019) and transposons integrated in the genomes of Deinococcus-Thermus, Bacteroidetes and Firmicutes hosts. The fourth clade represents transposons that are almost exclusively integrated into Proteobacteria hosts, including subtype I-F CASTs (Petassi et al., 2020), with only two branches containing Cyanobacteria and Firmicutes transposons. Additional hosts that are less common and were not reported in prior surveys for Tn7-like transposons (Parks and Peters, 2007, 2009) include Nitrospirae (8), Verrucomicrobia (5), Acidobacteria (4), Fibrobacteres (4), Aquificae (3), Spirochaetes (2), Deferribacteres (2) and Planctomycetes (1) (**Table S1**). No Tn7-like transposase homologs were identified in the currently available genomes of any archaea or viruses. Overall, the phylogenetic tree illustrates multiple switches between host phyla in the evolutionary history of Tn7-like transposons.

### Arrays of transposase binding sites reveal the boundaries of Tn7- like transposons

The transposons Tn7, Tn5090, Tn5053, and Tn552 possess arrays of inverted repeats located subterminal to the 5’ and 3’ boundaries of the transposon (Kamali-Moghaddam and Sundström, 2001), also known as the transposon’s left and right ends (LE/REs). The approximately 20 bp inverted repeats comprise the transposase binding sites and are typically located in intergenic regions (Arciszewska et al., 1989; Peters et al., 2017). We leveraged the presence of multiple inverted repeats to identify the transposon boundaries and discriminate intact transposons from partial ones. All intergenic sequences up to 125 kb from either side of *tnsB* were scanned for the presence of inverted repeat arrays resembling the LE/RE of Tn7 (see Methods). Under the premise that phylogenetically close transposases would utilize conserved binding site sequences, any discovered arrays of inverted repeats flanking a given transposase were compared to those found for another, closely related homolog, if available. If the inverted repeat sequences were conserved among ≥ 50% of close transposase homologs and satisfied additional criteria (see Methods), the arrays were predicted to be the LE/REs of the respective transposons. The computationally predicted TnsB binding sites were consistent with biochemical data on Tn7 transposition, where three TnsB binding sites were present at the left end and four at the right end (Arciszewska et al., 1989) (Fig 1B). Similar arrays were identified at the (putative) left and right ends of other transposons, despite the phylogenetic distance from Tn7 (Fig 1A). Thus, arrays of inverted repeats, which are candidate TnsB transposase binding sites, are a general feature that unites Tn7-like transposons.

To estimate the LE/RE detection accuracy, the predicted boundaries of 523 CASTs (Hsieh and Peters, 2021; Petassi et al., 2020) were analyzed. It is possible to identify the target site of the CASTs although transposition has not been demonstrated experimentally, by extracting spacers from the CRISPR array and searching the sequence neighborhood of the transposon for a matching protospacer. CASTs and other Tn7-like transposons integrate ∼50 bp downstream of the target site, where the 50 bp region between the target and integration site likely corresponds to the “footprint” of the transposition complex (Peters, 2015; Shen et al., 2021). In total, the LEs/REs were predicted for 352/523 (67%) of the previously reported CASTs. Almost all predicted integration sites are 50-60 bp from the protospacer-containing target site, with only two transposons predicted to be located more than 1 kb away from the attachment site (Fig. 1C). The 171 CASTs that lack both predicted ends could be incompletely sequenced (e.g., the transposon spans multiple contigs), have degenerate TnsB binding sites or LEs/REs that are otherwise undetectable by this approach and were not investigated further. The results of the CAST analysis indicate that the predicted TnsB binding sites represent the LE/REs of the transposons with substantial accuracy. Therefore, the genes embedded between the predicted TnsB binding sites in the LE/RE are not simply adjacent to *tnsB*, but rather, are mobilized by Tn7-like transposons.

#### Core gene content of Tn7-like transposons

Using the predicted TnsB binding sites to determine the outermost boundaries of the transposons, the nucleotide sequences were extracted from the respective contigs and dereplicated (99% average nucleotide identity across 95% alignment length), resulting in a final set of 6,549 putatively complete Tn7-like transposons. Besides the universally conserved *tnsB* transposase, which was a minimal requirement of the transposon discovery pipeline, 96% of the identified transposons harbored a *tnsC*-family ATPase (Fig 1D). The next most conserved gene is the target-site selector *tnsD/tniQ (*hereafter *tnsD*), which was present in 73-97% of the transposons in each clade. The PDEXK-family transposase *tnsA* is nearly ubiquitous in clade 1 (92%) but relatively uncommon in clade 3 (28%); this pattern is inverted for the tyrosine and serine superfamily resolvases (*tnsR)* (see detailed discussion in the following section). Only the transposons in clades 1 and 3 contained a homolog of *tnsE*, which promotes the integration of Tn7 into replication forks, including those on plasmids (Parks et al., 2009). A total of 1,298 transposons (20%) were found to be integrated into plasmids, including some lacking an identifiable *tnsE* homolog (**Table S2**). This discrepancy could be attributable to multiple factors, including incorrect assignment of a chromosomal contig as a plasmid, the inability to identify weakly-conserved *tnsE* homologs, the presence of a *trans-*acting TnsE, or involvement of other, as yet unknown plasmid target-site selecting genes. The median length of the transposons ranged from 14-29 kb (Fig. 1E), but much larger transposons were detected as well, the longest reaching 140 kb (**Table S2**). The cargo genes encoded within the boundaries of the transposons were functionally profiled, as described below.

### Tn7-like transposons cross horizontal gene transfer barriers

To complement the phylogenetic analysis and identify potential phylogenetic boundaries of horizontal transfer, a similarity matrix was constructed between all pairs of transposons using the number of shared protein clusters. Protein clusters were constructed greedily from ORFs, with the clustering threshold at 80% amino acid sequence identity over 75% of the length of the smaller ORF. Next, the similarity matrix between all transposons was analyzed to identify communities, that is, sets of transposons that shared genes significantly more frequently with each other than with transposons in the rest of the network. Using a hierarchical community-detection approach (Zheng et al., 2021), several large communities were identified at lower “resolutions” and then iteratively partitioned into smaller subcommunities at higher resolutions, until a reaching a predefined resolution limit. The communities that are stable across resolutions are considered significant, whereas transient communities are discarded, leveraging the network concept known as ‘persistent homology’ (Zheng et al., 2021). To select the predefined resolution limit, which constrains the size of the smallest subcommunities, a range of upper limits was explored and the limit that resulted in the highest mean community persistence (that is, quality) was chosen to identify the final communities of transposons (Fig. S1). In the resulting hierarchical depiction of the underlying community structure, the communities decrease in size and homogenize at the taxonomic level of phylum with descending levels of the hierarchy (Fig 2A). The phylum-level taxonomic homogeneity of the communities indicates that recent HGT via transposition between bacterial phyla is limited, in general. Nevertheless, a community of 1,221 promiscuous transposons that overcome this barrier was delineated. In this community, 83% of the transposons originated from Proteobacteria, and it could be divided into 15 subcommunities (Fig. 2B). Each subcommunity was labelled with the consensus last common ancestor, such that ≥ 80% of transposons in a subcommunity integrate into hosts that belong to the same taxonomic lineage. Inter-phyla exchange was found to occur between the *Micrococcales*-dominated subcommunity (Actinobacteria) and multiple classes of Proteobacteria, with an observable contribution of plasmid-mediated transfer (Fig. 2C). Altogether, for the 15 subcommunities, ≥80% of the transposons integrate into hosts from the same phylum but multiple classes (3 communities), the same order but multiple families (3 communities), the same family but multiple genera (5 communities) or the same genus but multiple species (2 communities); the remaining two subcommunities do not have a consensus phylogenetic level (Fig. 2B). We next tested if the communities that encompassed multiple phylogenetically distant hosts were due to a lack of resolution, that is, smaller subcommunities of closely related hosts were actually present but went undetected. Constructing more subcommunities of smaller size did not appreciably change the consensus phylogenetic level of the subcommunities at the lowest levels of the hierarchy (Fig. S1D), indicating that the analysis was not limited by resolution. Instead, the community composition apparently reflected promiscuous HGT between different genera, families, orders and classes of bacteria via transposons, along with sporadic exchanges between phyla.

**Figure 2.**
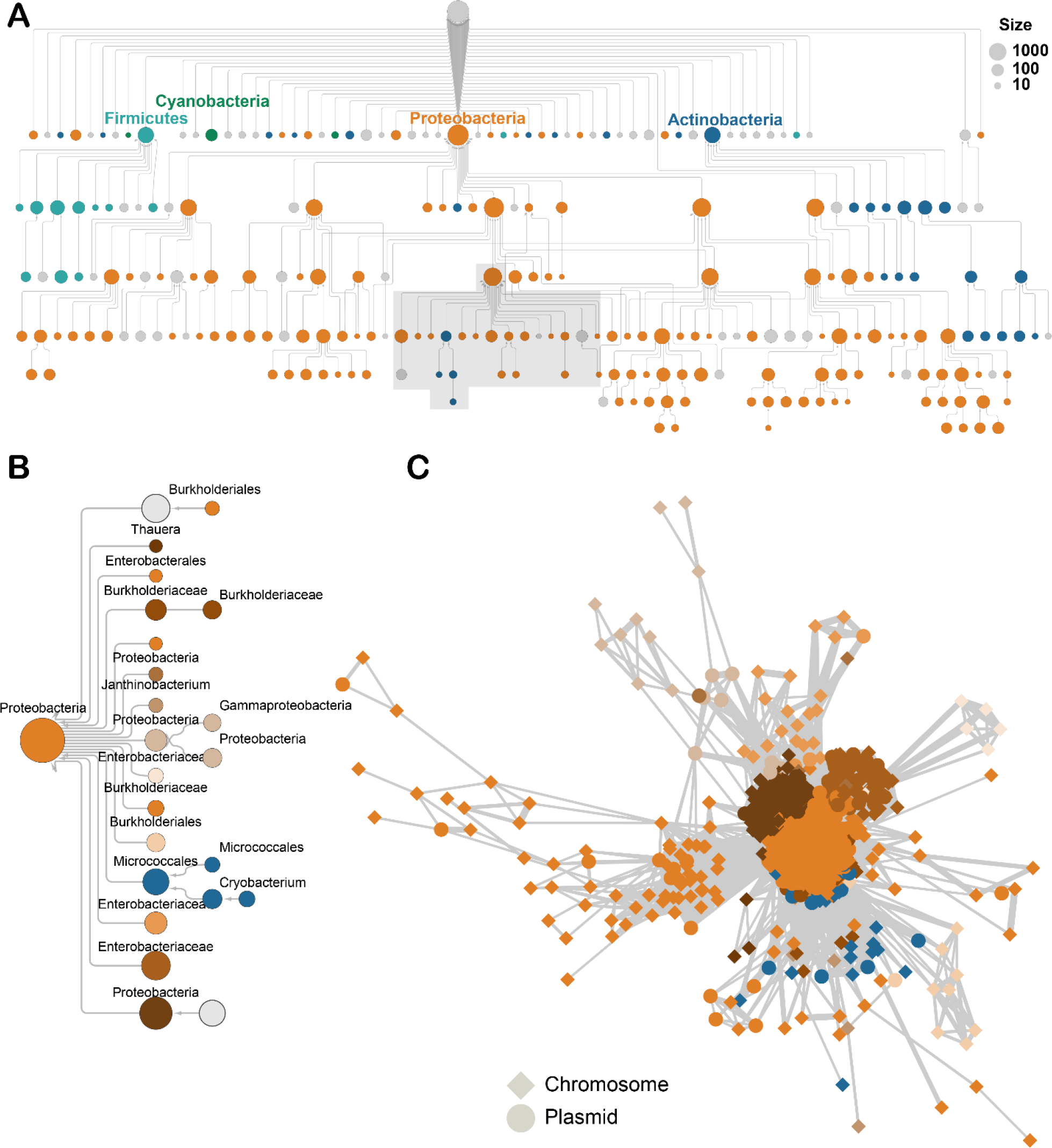
Tn7 transposons disseminate genes across deep phylogenetic divides. A hierarchical network of transposon communities (nodes), where each edge signifies the division of one community into another, smaller subcommunity. All communities are scaled by the total number of members and are colored if ≥ 80% of the members belong to the same phylum (**A**). Expansion of the area shaded in grey, with subcommunities recolored and labeled with their lowest common ancestor (80% consensus), if present (**B**). Network of 1,221 transposons (nodes) connected with edges that are weighted by the similarity between the respective two transposons. The shape of the node relates if the transposon is inserted into the host’s chromosome or a plasmid (**C**).

The transposon communities were further analyzed to determine if membership was driven by core or cargo genes. Pairs of transposons carrying identical cargo were common, including those from different clades. For example, two *Bacillus cereus* transposons from clades 1 and 2 showed minimal sequence similarity between their core proteins but both carry a nearly identical arsenate resistance operon and because of that belong to the same community (Fig. S2). Conversely, pairs of transposons in the same community could be found that mobilized entirely different cargo. For example, two transposons integrated in the same *Pseudomonas monteillii* genome shared no protein clusters, apart from the core proteins (Fig. S2), highlighting that closely related transposons can transport unrelated cargo. These contrasting examples demonstrate that community membership is defined by both core and cargo genes; furthermore, these findings emphasize that the cargo is decoupled from the transposition core machinery.

### Functional repertoire of Tn7 cargo includes other mobile genetic elements

To examine the functional repertoire of the cargo, the subset of the Tn7-like transposons found to be integrated into completely sequenced bacterial genomes (n = 739) were compared against randomly selected loci of equal lengths. All ORFs were annotated against the COG database (Galperin et al., 2020) and binned into the COG functional classes to estimate the relative frequency of each functional class in both gene sets. Obviously, homologs of the Tn7 core genes drive the enrichment of the mobilome COG class in transposons relative to random chromosomal loci (Fig. 3). However, other MGEs also contribute to the enrichment of the mobilome genes. The most common MGEs identified within Tn7-like transposons are insertion sequences (ISs), particularly those of the IS3 family, that are present in ∼30% of the transposons (Fig S3A). In some cases, two IS elements of the same family flank one or several genes from both sides within a Tn7-like transposon. Such proximity between two ISs of the same family can enable both to mobilize in tandem as a composite, cargo-carrying transposon (Siguier et al., 2015). Two examples of clinical importance are ISs that flank phosphoethanolamine transferases *mcr-9.1* or *mcr-3* (Fig S3B) that reduce the affinity of lipid A to colistin, a “last resort” antibiotic (Grégoire et al., 2017). The *mcr-3-*carrying transposon is nested in a Tn7-like transposon borne on a plasmid in *E. coli,* highlighting the broad mobile potential of these genes. A third example of potential ecological relevance was found in *Nitrospira moscoviensis,* where two IS21-family transposons flank cytochrome P460 genes that are involved in nitrogen cycling (Caranto et al., 2016), all within the boundaries of a Tn7-like transposon (Fig. S3C). Finally, resolvase-carrying transposons unrelated to Tn7 were also identified as cargo (Fig. S3D). Thus, one of the major sources of the Tn7-like transposon cargo are other MGEs, some of which carry their own cargo.

**Figure 3.**
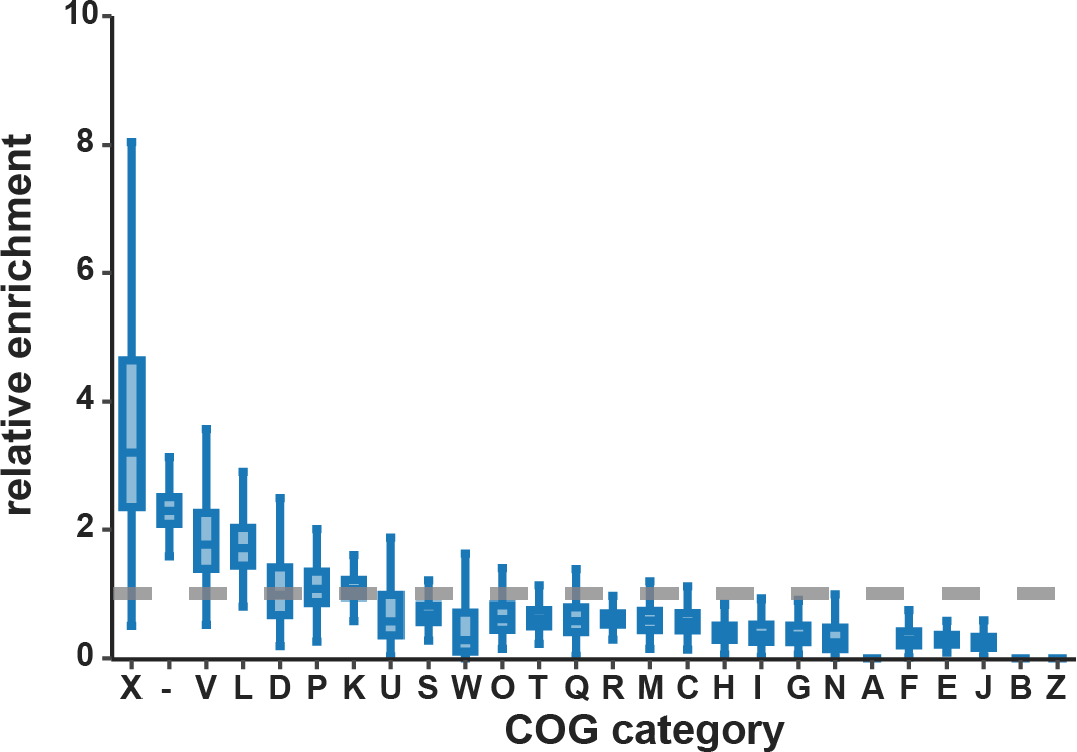
Tn7-like transposons mobilize a distinct repertoire of genes. A subset of transposons in completely sequenced bacteria (n = 739) were used to calculate the enrichment of individual COG categories in the transposons relative to randomly sampled genomic loci. The three enriched categories are “mobilome” (X), “defense” (V), and “replication, recombination, and repair” (L), as well as ORFs without significant similarity to a COG (“-“).

### Defense systems associated with Tn7-like transposons

The next most highly enriched set of genes, apart from genes of unknown function, comes from the defense COG class. Search of the cargo for genes involved in biological conflicts uncovered numerous, diverse innate immune systems, the most common and widespread being restriction-modification modules (Table 1). The next most frequent defense system includes the OLD-family nuclease, which contains a TOPRIM domain (Aravind et al., 1998) and interferes with the replication of phage Lambda (Sironi, 1969). The OLD nuclease is typically associated with a UvrD-family helicase (Fig. S4), jointly comprising the Gabija system that confers immunity against multiple phages (Doron et al., 2018). Another widely mobilized phage defense system is the cyclic oligonucleotide based antiphage signaling system (CBASS) (Millman et al., 2020), which includes a nucleotidyltransferase that recognizes phage proteins and mediates programmed cell death through various effectors (Cohen et al., 2019). Other, less abundant innate immune systems with diverse mechanisms were identified as well (Burroughs et al., 2013; Doron et al., 2018; Gao et al., 2020; Goldfarb et al., 2015; Wang et al., 2018) (Table 1). The transposons also mobilize single-component defense systems that confer protection against multiple dsDNA phages, including enzymes of the sirtuin superfamily (Gao et al., 2020), viperins (Bernheim et al., 2021), “defensive” reverse transcriptases (Gao et al., 2020) and enzymes that deplete the pool of nucleotides available for phage reproduction (Tal et al., 2021). Although these previously described innate immune systems jointly represent only 2% of the cargo genes, they are carried by 26% of the transposons. It appears likely that unknown defense systems are lurking among the cargo genes with currently unknown functions.

**Table 1:** Tn7-like transposons mobilize diverse immune systems. The number of transposons mobilizing each immune system is tabulated based on the listed components. The phyla are abbreviated as follows: Proteobacteria (Pro), Actinobacteria (Act), Firmicutes (Fir), Cyanobacteria (Cy), Chloroflexi (Chl), Spirochaetes (Spi), Bacteroides (Bac), Deinococcus-Thermus (DT), Nitrosomonas (Nit)

| Immune mechanism | System | Components | No. Tns | Phyla |
| --- | --- | --- | --- | --- |
| Innate | Restriction-modification | Restriction endonucleases (COG0610, COG4096, COG3440, COG1002, COG3421, COG1715, COG3587, COG4636, COG4804, COG1403, COG3183, COG4127, COG4748)<br>Methylases (COG0286), Specificity subunits (COG0732, COG4268, COG1401) | 947 | Pro, Act, Fir, Cy, Chl, Spi, Bac |
|  | Gabija | OLD nuclease (COG3593) | 338 | Pro, Fir, Act, Bac |
|  | Sirtuin | Sirtuin2 (cl00195) | 242 | Pro, Act, Fir, Cy |
|  | CBASS | Nucleotidyltransferase (cd05400) | 216 | Pro, Act, Fir |
|  | Thoeris | ThsB (pfam08937) | 61 | Pro, Act, Fir |
|  | Retron | RT (cd01646) | 50 | Pro, Act, Fir |
|  | Hachimann | HamA (pfam08878) | 43 | Pro, Fir, Spi |
|  | BREX | BrxC (NF033441) | 40 | Pro, Act, Fir, Cy, Nit, DT |
|  | Wadjet | JetD (COG4924) | 34 | Pro, Act |
|  | Kiwa | KwaB (pfam16162) | 23 | Pro, Act |
|  | Nucleotide depletion | dGTPase (PRK05318), dCTPase (cd01286) | 21 | Pro, Fir |
|  | PIWI | PIWI REase (pfam18154) | 19 | Pro, Act, Fir |
|  | DND | DndE (pfam08870), DndB (pfam14072) | 14 | Pro, Bac, Fir, Cy |
|  | Zorya | ZorA (pfam01618) | 3 | Pro |
|  | Viperin | Viperin (TIGR04278) | 1 | Pro |
| Toxin-Antitoxin | ImmA | ImmA (COG2856, pfam06114) | 295 | Pro, Act, Bac, Fir, Cy |
|  | PIN | PIN (cl28905) | 251 | Pro, Act, Fir, Cy |
|  | ParE | ParE (cl21503) | 120 | Pro, Act, Cy, DT |
|  | RES | Res (cl02411) | 95 | Pro, Act, Fir |
|  | RelA-SpoT | RelA (cd05403, COG1669, smart00954, cd05399, pfam14907, COG2357, pfam04607, pfam01909, cd07749) | 70 | Pro, Act, Bac, Fir, Cy |
|  | HipA | HipA (cl28916) | 51 | Pro, Act, Cy |
|  | AbiC | AbiC (pfam14355) | 48 | Pro, Fir |
|  | HEPN | HEPN (cl00824) | 37 | Pro, Fir, Cy |
|  | AbiF | AbiF (pfam07751, COG4823) | 23 | Pro, Spi |
|  | AbiE | AbiEii (pfam08843, COG4849) | 19 | Pro, Act |
|  | Fic/Doc | Doc (COG3654, TIGR02613) | 7 | Pro, Cy |
|  | Barnase | BrnT (pfam04365, COG2929) | 5 | Pro, Cy |
| Adaptive | minimal I-F | Cas6_I-F, Csy3_I-F, Csy2_I-F | 304 | Pro |
|  | minimal V-K | Cas12k | 78 | Cy |
|  | minimal I-B | Multiple | 9 | Cy |
|  | Type III | Cas10 | 2 | Pro |
|  | Type I-B | Multiple | 1 | Cy |

Mobile genetic elements often encompass toxin-antitoxin (TA) gene pairs, which frequently colocalize with innate immune systems (Makarova et al., 2009, 2013) and can function as defense systems themselves, typically eliciting dormancy or programmed cell death (Chopin et al., 2005; Harms et al., 2018; Page and Peti, 2016). Tn7-like transposons mobilize a variety of toxin effectors, including PIN family ribonucleases, RelA/SpoT-like nucleotidyltransferases, ParE-family mRNA interferases, ADP-ribosyltransferases and HipA-family kinases (**Table 1**). Several abortive infection (AI) systems first described in *Lactococci* were also identified (Labrie and Moineau, 2007), implicating transposons in horizontal transfer of these genes. In multiple phyla, the toxins are associated with a cognate antitoxin and/or a defense system (Fig. S5). Thus, TAs and the functionally similar AI systems substantially augment the repertoire of innate immune systems carried by Tn7-like transposons.

To identify transposons that mobilize adaptive immune systems, CRISPR arrays were predicted and the cargo genes were scanned for significant sequence similarity to *cas* genes. In this survey, 410 transposons were shown to harbor at least one *cas* gene or a CRISPR array; of these, all but 14 belong to one of three clades in the TnsB phylogenetic tree (Fig. S6). The location of CASTs in the TnsB tree demonstrates that Tn7-like transposons exapted CRISPR-Cas systems for target site selection on at least three independent occasions, in agreement with previous observations (Faure et al., 2019; Peters et al., 2017). The LE/REs of one experimentally characterized CAST (CAST I-B2) (Saito et al., 2021) could not be predicted automatically, raising the possibility that improvements in transposon delineation could reveal novel CASTs. Additional transposons scattered across the TnsB tree contain individual components of CRISPR-Cas systems or, less frequently, a complete system (Table 1, Fig. S6). Specifically, one *Geminocystis spp.* transposon is closely related to subtype I-B CASTs but carries *cas2* next to a CRISPR array and a *cas1-cas4* fusion that together comprise the adaptation module, which is normally absent in CASTs (Fig. S6). Interestingly, this bacterium also hosts two subtype V-K CASTs which are integrated at different loci. Two other transposons encode a type III CRISPR-Cas system with the hallmark *cas10* effector (Fig. S6). Overall, Tn7-like transposons infrequently possess a complete, fully functional adaptive immune system, but often carry a minimal suite of CRISPR-Cas genes that functions as the target site selection machinery.

### The replicative transposition pathway of Tn7-like transposons is predicted by the loss of a cut-and-paste transposase and gain of a resolvase

The final COG class that is enriched in the Tn7-like transposon cargo is “replication, recombination and repair”, and more specifically, genes encoding enzymes of the tyrosine and serine integrase/resolvase superfamilies. Enzymes from both superfamilies are encoded by diverse MGEs and catalyze site-specific rearrangements in DNA that are involved in transposon relocation from a donor to an acceptor site (Craig, 2015). For example, Tn3 and Tn21 transposition involves a cointegrate intermediate that is resolved into two separate molecules by a serine and a tyrosine superfamily resolvase, respectively (Nicolas et al., 2015). Because resolvases can be functionally involved in MGE mobility, they neither fit the strict definition of “cargo” nor can be clearly classified into the “replication, recombination and repair” or “mobilome” COG functional categories. Therefore, the genetic context and phyletic distribution of the serine and tyrosine superfamily enzymes were examined separately to better ascertain their functional roles.

In the collection of Tn7-like transposons analyzed here, serine and tyrosine resolvases are prevalent, but not universally conserved, in clades 2 and 3. Clade 3 includes Tn5053 and other transposons that possess either a serine or a tyrosine resolvase, whereas clade 2 transposons are almost exclusively associated with tyrosine resolvases (Fig. S7). Both these clades contain branches of transposons that lack homologs of *tnsA*, which is a transposase essential for cut-and-paste transposition in Tn7 (Peters, 2015) (Fig. 1, Fig. S7). The absence of *tnsA* combined with the presence of a serine or a tyrosine resolvase suggests that these transposons do not mobilize via the cut-and-paste mechanism. Instead, their transposition likely proceeds through a cointegrate intermediate that needs to be resolved prior to integration. Transposons that lack a cognate resolvase can still mobilize via a cointegrate intermediate, where resolution is achieved by resolvase *in trans* or by the host *recA*-mediated homologous recombination (May and Craig, 1996; Minakhina et al., 1999; Nicolas et al., 2015), perhaps explaining the lack of resolvases in some of these transposons. Together, these findings predict that replicative transposition is the principal pathway employed by distinct branches of transposons within these two clades, in which case the resolvases are not cargo but rather core components of the transposition machinery.

Numerous other transposons outside of these two branches also harbor serine resolvases, but their scatter across the TnsB tree and presence of *tnsA* homologs does not support a functional role during replicative transposition (Fig. S7). In some cases, the serine resolvase is encoded as part of a nested transposon (Fig. S3C). Here, the resolvase is a core component of the Tn3-family transposon, but appears to be cargo for the larger, Tn7-like transposon, although it cannot be ruled out that the resolvase functions during replicative transposition of both transposons. In other cases, the serine resolvase does not appear to be part of a nested transposon, suggesting it is only carried as cargo by the Tn7-like transposon and might perform other functions, such as DNA inversion.

### Cargo genes involved in antibiotic-resistance, biosynthesis and central carbon metabolism

To characterize the cargo genes that belong to COG functional classes that are not enriched or even are depleted (Fig. 3), the specific metabolic or functional pathways were tabulated for each individual COG. Few complete pathways were carried by any of the Tn7-like transposons, but 37% of the transposons harbor at least one gene implicated in various aspects of cellular physiology, ranging from amino acid catabolism to nucleotide and lipid biosynthesis (Fig. 4). The most common, albeit incomplete, pathway is folate biosynthesis (n = 555 transposons, category “H”), which includes the genes *sul1* and *dfrA1* encoding, respectively, dihydropteroate synthase and dihydrofolate reductase which confer resistance to sulfonamide antibiotics (Huovinen et al., 1995). Similarly, the mercury resistance gene *merA* (COG1249, category “E”) is responsible for the apparent commonality of the glycine cleavage pathway. To explore the larger pool of antibiotic resistance genes, which belong to various COG categories, the cargo was searched against a database of genes with experimentally demonstrated roles in antibiotic and xenobiotic resistance (Feldgarden et al., 2019). Genes with heavy metal detoxification activity, including mercury, were found to be abundant, as well as antibiotic resistance genes that confer protection against the aminoglycoside, sulfonamide and beta-lactam classes of antibiotics (Fig. S8). Thus, Tn7-like transposons commonly mobilize genes of biological and clinical relevance.

**Figure 4:**
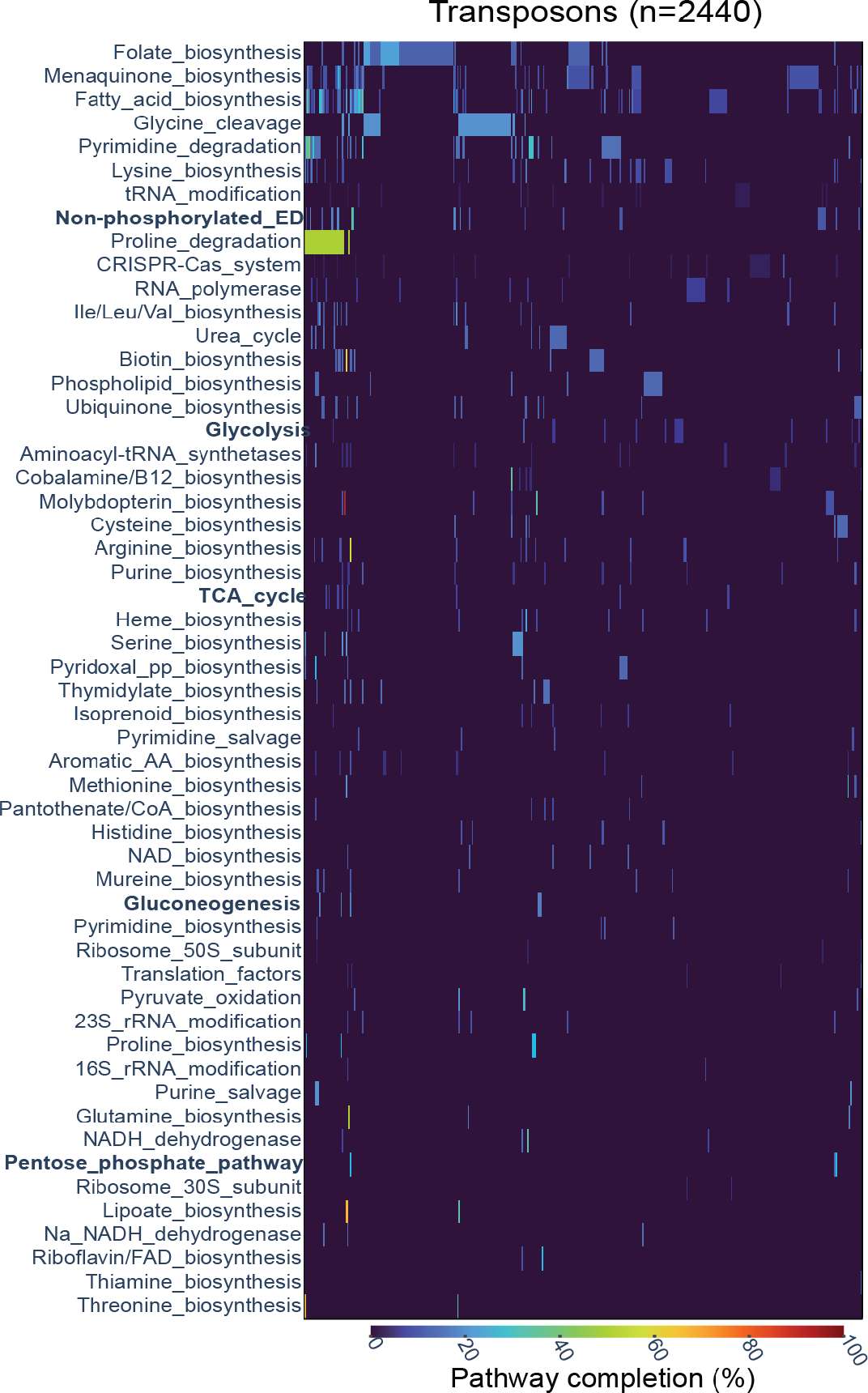
Metabolic pathways mobilized by Tn7-like transposons. The x-axis lists the COG pathways, sorted by the pathways that are found in the most transposons to least. Each column in the y-axis corresponds to an individual transposon, with the color of the cell proportional to the completeness of the individual pathway. Central carbon metabolism pathways discussed in the main text are in bold.

To elucidate how antibiotic and stress-resistance genes are captured by transposons, the cargo was examined for the presence of integrons. Integrons are MGE that site-specifically incorporate “cassettes” of DNA and comprise a notable component of the cargo of Tn7 and other families of transposons because of their role in aggregating antibiotic resistance genes (Escudero et al., 2015; Ramírez et al., 2005). Altogether, 10% of the Tn7-like transposons harbor an intact or partial integron, collectively encoding 2334 integron gene cassettes (i.e., ORFs with a 3’ *attC* site). The cassettes are dominated by *qacE* (n = 334 cassettes) and *dfrA1* (n = 157 cassettes) that confer resistance to ammonium-based disinfectants and trimethoprim, respectively. Integrons carrying homologs of genes that provide resistance to aminoglycosides, beta-lactams, phenicols, sulfonamides, macrolides, lincosamides, quinolones and bleomycins were also captured by Tn7- like transposons, illuminating the role of integrons in expanding the diversity of the antibiotic- and stress-resistance cargo.

The transposons mobilizing genes involved in biosynthetic pathways and central carbon metabolism were analyzed next. The most common biosynthetic pathways in transposons are those for fatty acids and menaquinones, including COG1028, short-chain alcohol dehydrogenase, COG1960, acyl-CoA dehydrogenase, COG0183, acetyl-coA acetyltransferase, and COG0596, MenH-family esterase. These enzymes appear in a variety of genetic contexts, making their specific roles and end products difficult to infer, but generally are involved in the synthesis or degradation of “auxiliary” compounds. Additionally, multiple transposons carry genes involved in central metabolic processes, including glycolysis, gluconeogenesis, pentose phosphate pathway and tricarboxylic acid cycle (Fig. 4). The genetic context of these central carbon metabolism genes includes other genes involved in sugar catabolism and transport (Fig. S9). Overall, although the components of core and auxiliary metabolic pathways are not common cargo compared to their representation in bacterial genomes, their presence broadly reveals the potential for Tn7-like transposons to mobilize genes that modulate cellular metabolism.

## Discussion

A search of prokaryotic genomes initiated with HMMs for the core Tn7 proteins involved in transposition recovered thousands of predicted transposons in diverse bacterial phyla, underscoring the mobility of these elements. The phylogeny of TnsB demonstrates the relationship between Tn7 and other well-characterized transposons, including Tn5053 and the recently described CASTs. Besides the phylogenetic coherence of the transposase, all these transposons possess similar architectures of inverted repeats at their left and right ends that mark the boundaries of the transposon. Moreover, within these boundaries, most encode a transposition-regulating ATPase (*tnsC*) and a target site-selecting protein (*tnsD)* flanking the core transposase. These shared features are the hallmarks of a vast, distinct group of MGE informally dubbed “Tn7-like” transposons (Oppon et al., 1998; Parks and Peters, 2009)

Tn7-like transposons encode a plastic repertoire of genes, even among the genes that are directly involved in transposition. A prominent example of such plasticity is *tnsA*, where point mutations in the catalytic site abolish cut-and-paste transposition but do not impede replicative transposition (May and Craig, 1996; Vo et al., 2021). Multiple branches of Tn7-like transposons lack *tnsA* altogether (Fig. S7), evincing the dispensability of this gene. Typically, the *tnsA-* lacking transposons gain a tyrosine- or serine-superfamily resolvase (*tnsR)* that most likely mediates replicative transposition. The target site-selecting gene *tnsE* is absent in most clades of transposons although the actual spread of this gene might be underestimated due to the weak sequence conservation. Similarly, the CRISPR-Cas target site-selecting systems are relatively rare, but additional CASTs might be discovered with improvements in transposon delineation. The evolutionary gain and loss of *tnsAER* and the specialized CRISPR-Cas systems illustrate the modularity of the transposition machinery of the Tn7-like transposons.

The cargo genes carried by Tn7-like transposons exhibit an even greater plasticity. To survey the biological functions of the cargo, a comparison against genes that are encoded outside of the transposon was undertaken. An unexpected outcome of this analysis was that the most highly enriched function, the “mobilome”, was not explained by the presence of core Tn7 proteins alone but also by substantial contributions of other MGEs, specifically, IS elements. The enrichment of ISs suggests a stepwise process of gene capture by Tn7-like transposons, beginning with the formation of a compound transposon by two ISs at a locus outside of the Tn7- like element, followed the compound transposon jumping into the boundaries of the element, yielding a “nested” transposon. Such “nesting” of MGEs has been observed in the analysis of individual transposons and other MGEs (Krupovic et al., 2019; Liebert et al., 1999; Minakhina et al., 1999; Peters et al., 2014; Sota et al., 2002). The results presented here confirm that MGEs are a widespread, prominent source of the cargo carried by Tn7-like transposons. Accretion of MGEs apparently complements other mechanisms proposed to be involved in the capture of genes by transposons, such as homologous recombination (Nicolas et al., 2015; Partridge, 2011), culminating in the diverse and seemingly haphazard repertoire of cargo genes observed here. The relative contribution of MGE accretion versus homologous recombination to the diversity of cargo was not addressed here, but this question is now tractable with the database of transposons compiled in this and other studies (Ross et al., 2021).

One of the principal forces limiting HGT in prokaryotes is the cost of assimilating new genes into existing, regulated networks (Bershtein et al., 2015; Iranzo et al., 2017; Sela et al., 2019; Sorek et al., 2007). Integration itself can result in fitness consequences for the host cell, for example by disrupting and hence inactivating a gene (Durrant et al., 2020). Tn7-like transposons appear to integrate into “safe sites”, such as downstream of tRNAs, using dedicated target site-selecting proteins, leaving the attachment site intact and thus preempting a potential deleterious effect for the host cell (Peters, 2019). Beyond the consequences of integration itself, the introduction of horizontally acquired genes into an existing network can affect host fitness. A case in point are the antibiotic resistance genes mobilized by hundreds of Tn7-like transposons identified in this study, many of which were captured via integrons. The additional burden of the integron cassettes is offset by endowing the cell to survive in the presence of antibiotics (Liebert et al., 1999). Analogously, a similar fraction of the Tn7-like transposons mobilize a diverse compendium of genes that confer resistance to bacteriophages (**Table 1**). Although possession of defense systems comes at a high cost due to autoimmunity and other effects (Cui and Bikard, 2016; Heussler et al., 2015; Iranzo et al., 2017; Koonin and Zhang, 2017; Pleška et al., 2016), their enrichment in Tn7-like transposons implies that the cost is superseded by the benefit of protection against phage predation. The concurrent circulation of antibiotic and phage resistance genes is also observed in other MGEs (LeGault et al., 2020), highlighting how the cargo of a single MGE can confer immunity to both agents and redresses the cost of harmonization with the existing cellular network. An additional twist on the subject of defense systems in Tn7-like transposons is the incorporation of CRISPR arrays that are apparently involved in competition with other MGEs (Faure et al., 2019).

The metabolic cargo genes carried by Tn7-like transposons offer a unique perspective into the energetic demands of their hosts. Intuitively, these genes would reflect the diverse niches occupied by each host, which span a variety of habitats, from the open ocean to the human gut. For example, a *Nitrospira moscoviensis* Tn7-like transposon carries a P460 cytochrome involved in the oxidation of hydroxylamine to nitrous oxide (Caranto et al., 2016), divulging the importance of the ammonia oxidation pathway for these soil-dwelling bacteria. The widespread transfer of *sul1* and *dfrA1* involved in the synthesis of folic acid is likely driven by environmental exposure to antibiotics that target this pathway. Transposons in the genera *Vibrio, Marinobacter, Bacillus, Rhizobia* and others harbor genes that metabolize compounds involved in central carbon metabolism pathways, again prompting the question of how these “core” metabolic genes harmonize with the existing network upon introduction into a new host genome. The identification of these cargo-carrying transposons will facilitate experimental efforts aimed at this question.

The Tn7-like transposons encode an expansive repertoire of genes that are functionally distinct from the rest of the host genome, raising the question how certain genes are captured and maintained whereas others are either never captured or are purged from the elements. The evidence presented here implies that any gene can, at one point, be encoded within a transposon, if only ephemerally. Mechanistically, this notion is supported by the prominence of ISs and compound transposons that vectorize genes (Siguier et al., 2015; Toleman et al., 2006) into Tn7- like transposons (Fig. S3), conceivably providing the transposon access to any gene in the host genome. Under this premise, the question becomes what selective forces result in the non-uniform distribution of the functions of cargo genes (Fig. 3). On one hand, anti-phage and antibiotic resistance genes appear to enhance the fitness of the host cell and favor their retention in the transposons. Furthermore, these genes as well as TA modules can make the host cell addicted to the transposons such that loss of a transposon results in cell death. On the other hand, the observation that certain functions are rarely represented in the cargo (for example, information processing, COG category “J”) implies that their continual presence in transposons is unfavorable to the host and hence to the transposons. Selection against these genes might manifest for a variety of reasons and drive their loss from the transposons. For example, and in particular, in the case of translation system components, expression of the respective genes from a transposon might result in a deleterious disruption of the stoichiometry of the protein complexes involved in these processes (Veitia, 2004; Veitia and Potier, 2015).

## Methods

### Identification of Tn7-like transposons

Multiple sequence alignments (MSAs) of the five core genes from *E. coli* Tn7 (*tnsABCDE)* were collected from the Pfam database using the following accessions: TnsA, PF08721 and PF08722; TnsB: PF00665; TnsC: PF11426 and PF05621; TnsD: PF15978; TniQ: PF06527; TnsE: PF18623. Additional MSAs of Tn7-like core proteins were generated from CASTs, and all MSAs were converted to Hidden Markov Models (HMMs) with HMMer (v. 3.1b2). For the purpose of a comprehensive search for Tn7-like transposons, a nucleotide database encompassing the NCBI WGS database and the MG-RAST database of metagenomes (Wilke et al., 2015) was prepared. The nucleotide sequences were input to Prodigal (v. 2.6.3) (Hyatt et al., 2010) for open reading frame (ORF) prediction. The collection of ORFs was queried with the Tn7 HMMs, using model-specific gathering cutoffs for the Pfam HMMs and the following cutoffs for the custom HMMs: -T 25 --domT 25 --incT 25 --incdomT 25. Any ORFs that produced a hit but were located less than 3 kb from the contig boundary were discarded to remove incomplete transposons. Additional heuristics were employed to select candidate Tn7- like transposons, using the following criteria: (1) a hit to the TnsB HMM with a bitscore ≥ 60, (2) a hit to at least one other Tn7 HMM and (3) two of the hits are in a putative operon, which is operationally defined as two codirected ORFs separated by less than 50 bp of non-coding sequence. All loci satisfying these criteria were considered candidate Tn7-like transposons and the putative TnsB orthologs were subjected to phylogenetic analysis.

The taxonomic information for each contig harboring a Tn7-like transposon was extracted from the NCBI taxonomy database using the entrez suite of command line tools. If a sequence was not indexed in the database, it was assigned to the domain Bacteria, given that Tn7-like transposons have not been so far identified in genomes of archaea or viruses (see results). Contigs were scanned using ViralVerify (Antipov et al., 2020) and the default database of HMMs to classify contigs as to viral, chromosomal, or plasmids.

#### Phylogenetic analysis

A phylogenetic tree of the DDE-family TnsB transposase was constructed using a previously described approach (Wolf et al., 2018). Briefly, the transposase ORFs were first clustered at 80% amino acid sequence identity over 75% of the length of the shorter ORF, and then, the representative sequences were re-clustered at 50% identity using MMSeqs2 (v. 12- 113e3) (Steinegger and Söding, 2018). The secondary cluster members were aligned using MUSCLE (Edgar, 2004) and compared to one another using HHSearch (Steinegger et al., 2019). The HHSearch similarity scores were used to construct an unweighted pair group method with arithmetic mean (UPGMA) dendrogram. The dendrogram was used to guide the pairwise alignment of clusters with HHalign (Steinegger et al., 2019). Any clusters that could not be aligned using this approach were discarded. The single resulting alignment was filtered to remove partial sequences and sites with more than 50% gaps and homogeneity lower than 0.1 (Yutin et al., 2008). An approximate maximum likelihood tree was constructed from the filtered alignment using FastTree2 (Price et al., 2010) with the Whelan-Goldman models of amino acid evolution and gamma-distributed site rates. Next, the tree was ultrametricized and branches were collapsed if the phylogenetic distance between them was less than 1. A single representative sequence was selected arbitrarily from each collapsed branch. The representatives were extracted from the main alignment and input to IQ-Tree (Nguyen et al., 2014) for phylogenetic reconstruction with parameters set to perform the aBayes branch test (Anisimova et al., 2011), ultrafast bootstrap approximation (Hoang et al., 2017) and automatic model selection (Kalyaanamoorthy et al., 2017), which selected LG+F+R10 as the best model. The tree was visualized using the interactive tree of life (Letunic and Bork, 2021).

### Delineating Tn7-like transposon boundaries

For a contig harboring a Tn7-like transposon, all intergenic sequences longer than 50 bp were extracted into a single fasta-formatted file. Coding sequences were excluded given that the boundaries of most Tn7-like transposons do not overlap with a predicted ORF (Faure et al., 2019; Peters et al., 2017). The intergenic sequences were used to construct a Markov model of the AT/GC content of the contig using the “fasta-get-markov” script provided in the MEME suite of tools (v. 5.3.0) (Bailey et al., 2009). The Markov model was used as a background file for all motif detection steps described in the following section. As Tn7-like transposons up to 117 kb in length have been reported (Peters et al., 2017), all intergenic sequences up to 125 kb on either side of a putative *tnsB* homolog were collected and stored in a separate file. An all-versus-all BLASTn search was executed with the following parameters: -word_size 4 -max_hsps 100 –evalue 100. The left and right ends of *E. coli* Tn7 contain 22 bp inverted repeats (Arciszewska et al., 1989), so the output was filtered for inverted repeats 15-40 bp in length, with no more than two gaps and 6 mismatches. Two intergenic sequences were considered candidate left and right ends if at least one of these sequences was located less than 20 kb from the start codon of the *tnsB* transposase and the two sequences shared two or more inverted repeats. Each unique combination of two intergenic sequences satisfying these criteria were input to MEME to detect nucleotide motifs.

The motif discovery was initiated with the following command line options: `-mod anr - nmotifs 10 -minw 15 -maxw 20 -minsites 4 –maxsites 6 -revcomp -markov_order 0 -evt 0.1`. This combination of parameters was selected on the basis that Tn7 encodes three TnsB binding sites at the left end and four at the right end (Arciszewska et al., 1989). The output was filtered to remove motifs with p-values greater that 10^-5^, present in only one of the two input sequences, and/or all located on the same strand. All motifs from all pairs of sequences that passed these filters were collected. The resulting set of motifs, representing putative TnsB binding sites, was used to search the 125 kb window of intergenic sequences around *tnsB* using the program FIMO (Grant et al., 2011), constrained with a 0.1 false discovery rate (q-value). The same filtering approach for the BLASTn output described above was applied to the FIMO output, with the following additional criteria: (1) the spacing between any two instances of a motif be ≤ 75 bp, (2) the total length of all motif instances on a sequence is ≤ 120 bp, (3) the motif cannot be present on 5 or more intergenic sequences, and (4) the product of the motif’s estimated false-discovery rate (combined q-value) be less than 0.01. Criterion 1 and 2 were enforced to match the spacing and combined length of TnsB binding sites in Tn7 (Arciszewska et al., 1989), whereas criterion 4 and 5 were applied to remove common, weakly-conserved nucleotide motifs. The single motif with the lowest combined q-value satisfying all of these criteria was selected as the motif that best represents the binding site of the respective input transposase.

The individual motifs from closely related *tnsB* homologs were next compared to one another. The motifs were compiled into a single file if the phylogenetic distance between the respective transposases was less than 1 (see the preceding section for details on the construction of the TnsB tree). Each file was input individually to TomTom (Gupta et al., 2007) for an all-versus-all motif alignment, using a minimum alignment length of 10 and the distance metric set as ‘pearson’. Any motifs that did not align with ≥ 50% of the motifs present in the file were discarded; otherwise, they were collected into a single, non-redundant motif dataset. All of the TnsB-centered windows of intergenic sequences were subjected to a final, competitive search against the motif dataset using FIMO. The best motif was selected using the FIMO filtering criteria described above and the outermost nucleotide coordinates of the motifs were recorded as the boundaries of the transposon.

### Annotation of genes encoded by Tn7-like transposons

The nucleotide sequence of each transposon was extracted from the respective contig using the predicted left and right boundaries. The ORFs were predicted using Prodigal in metagenomic mode (v. 2.6.3) (Hyatt et al., 2010) and clustered at 80% amino acid identity across 75% of the length of the shorter ORF using MMseqs2 (v. 12-113e3) (Steinegger and Söding, 2018). Multiple sequence alignments (MSAs) from the NCBI conserved domain database (v. 3.19) (Lu et al., 2019) database were used to query the representative ORFs of each cluster using PSI-BLAST (Altschul et al., 1997) with a 0.01 e-value cutoff. The ORFs that did not produce a significant PSI-BLAST match were subjected to an additional round of annotation. Each of these ORFs was queried against the Uniprot database clustered at 30% identity (constructed in June 2020, available at http://wwwuser.gwdg.de/∼compbiol/uniclust/) with HHblits (Steinegger et al., 2019), enforcing that any database sequence aligned with ≥ 20% of the query. The resulting alignments were used for a second iteration of the search and/or terminated if the number of effective sequences in the alignment was greater than 10. The MSAs were then used to query HHsuite-formatted databases of alignments from the PDB and NCBI conserved domain database, accepting hits with probability greater than 90. Any annotations assigned to the representative ORF of a given cluster were transferred to each member of the cluster.

The annotations obtained as described above were revised in the following cases to be consistent with field-specific nomenclature. For experimentally characterized antibiotic resistance genes, representative ORFs were scanned using AMRFinderPlus (v. 3.10.15) (Feldgarden et al., 2019) with default settings. For *cas* genes, representative ORFs were searched using PSI-BLAST against a database of multiple sequence alignments obtained from a recent survey (Makarova et al., 2020b). For insertion sequences, the annotations were manually updated to the family-level designation according to the ISFinder database (Siguier et al., 2006).

Non-coding RNAs were annotated using Infernal (v. 1.1.3) (Nawrocki and Eddy, 2013) against the Rfam database (v. 14.5) (Kalvari et al., 2020) using the model-specific bit score gathering threshold as a cutoff for significance. CRISPR arrays were detected using Minced (v. 0.4.2, https://github.com/ctSkennerton/minced) with the default settings. Integron *attC* sites were annotated using IntegronFinder (v. 2.0) (Cury et al., 2016). Annotations were displayed graphically using custom scripts and Clinker (v. 0.0.21) (Gilchrist and Chooi, 2021).

### Transposon dereplication, gene sharing network and community detection

The nucleotide sequences of the transposons were dereplicated at 99% average nucleotide identity across 95% of the contig length using dRep (Olm et al., 2017) and associated dependencies (Jain et al., 2018; Ondov et al., 2016). The protein clusters from dereplicated transposons were used to calculate a similarity matrix as previously described (Makarova et al., 2020a). The similarity matrix was input to Hidef (v. 1.0.0) (Zheng et al., 2021) to construct a hierarchical network of gene-sharing communities. Hidef requires a maximum resolution limit, which dictates the total number of communities and their size. To optimize this parameter, the similarity matrix was scanned with the maximum resolution parameter incremented in steps of 5, up to 1000. At each step, the output was parsed to assign transposons to their smallest community. The total number, size, persistence and Last Common Ancestor (LCA) of each community was tabulated. The LCA was defined using an 80% consensus rule, such that ≥ 80% of the transposons in a community have the same ancestor. A maximum resolution size of 140 yielded the highest mean community persistence without major changes to the phylogenetic level of the LCA (see Results), so the final network was constructed using this resolution and a persistence cutoff of 20; all other parameters were left as default. The LCA of each community in the final network was retabulated using the same 80% consensus rule, but without assigning transposons to their single, smallest subcommunity, so that the taxonomic makeup of communities at all hierarchical levels could be determined. Networks were visualized using Cytoscape (v. 3.7.2) (Shannon et al., 2003).

### Functional annotation of transposon genes using COGs

The enrichment or depletion of COG functional categories for the subset of transposons in completely sequenced bacterial genomes were calculated as described previously (Shmakov et al., 2020). Briefly, the ORFs encoded by these genomes were annotated by comparison to the COG database of multiple sequence alignments (Galperin et al., 2020) using the same PSI-BLAST parameters as described above. The COG categories corresponding to the transposon-encoded ORFs, or a randomly selected genomic locus of identical length, were extracted and summed. The relative frequency of each COG category in the transposons versus the genomic loci was calculated from a random selection of 100 pairs of transposons and their genomic equivalents, iterated 1000 times.

The COG annotations of transposon-encoded ORFs were extracted and assigned to their respective COG pathways. The completeness of the pathway was calculated as the number of COGs from the pathway represented in the transposon divided by the total number of COGs constituting the pathway. Pathways were discarded if represented in fewer than 10 transposons.

## Data availability

All sequences are publicly available from the NCBI and MG-RAST databases. The nucleotide coordinates of all transposons are provided in Table S2. Fasta-formatted files, annotations for all transposon open reading frames and source data for figures 1–3 are available via (https://ftp.ncbi.nih.gov/pub/yutinn/benler_2021/Tn7/source_data/).

---

**Figure S1.**
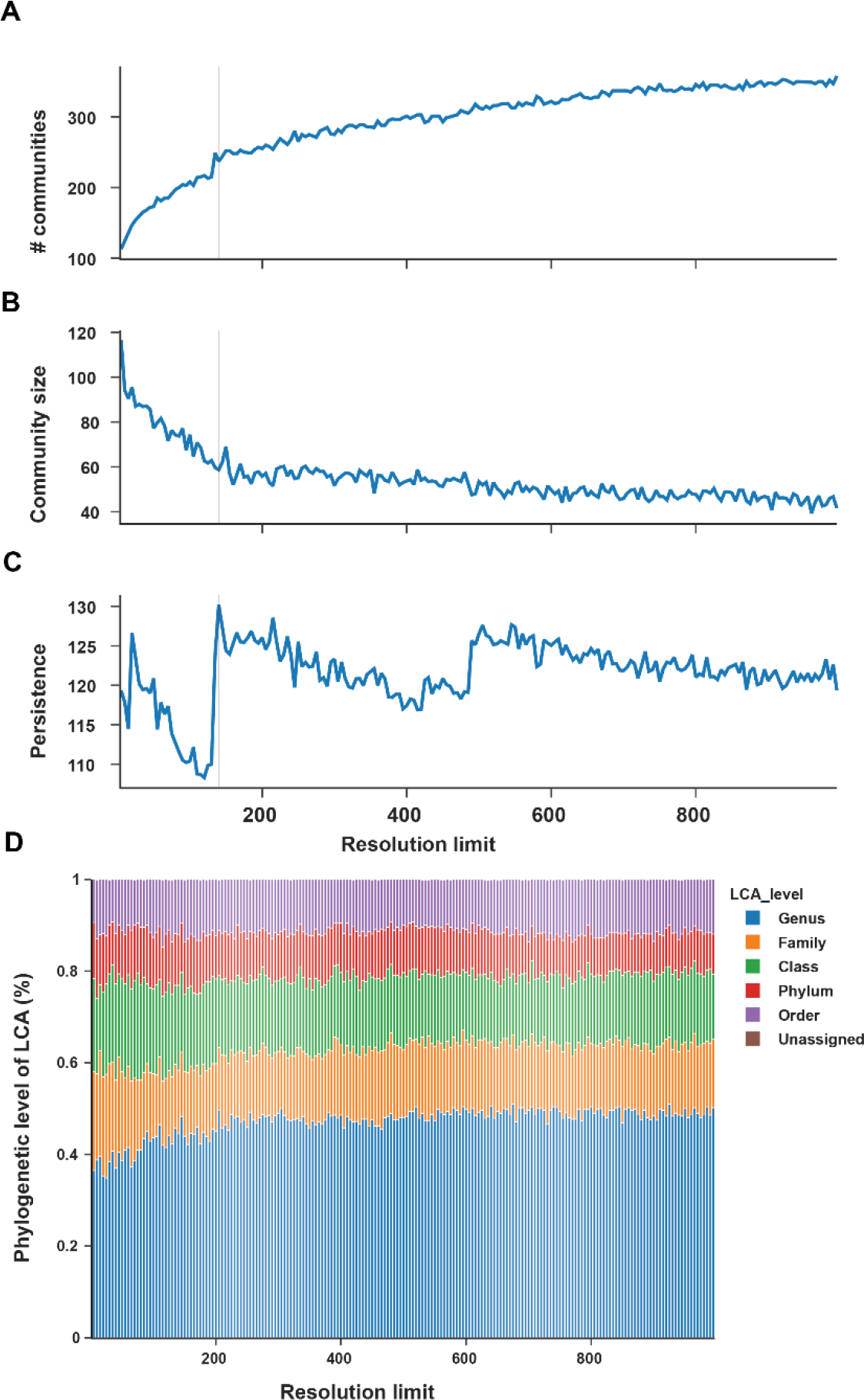
Selection of the resolution parameter to construct transposon communities. Transposon communities were detected given “resolution” limits between 5 and 1000, in steps of 5. Increasing the resolution leads to more communities (**A**) that are smaller, on average (**B**). A resolution of 140 yields the highest mean community persistence and was therefore selected to construct the final network (**C**). Construction of smaller communities marginally decreases the phylogenetic level of the communities’ lowest common ancestor, operationally defined as the consensus host of ≥ 80% of the transposons in a community (**D**).

**Figure S2.**
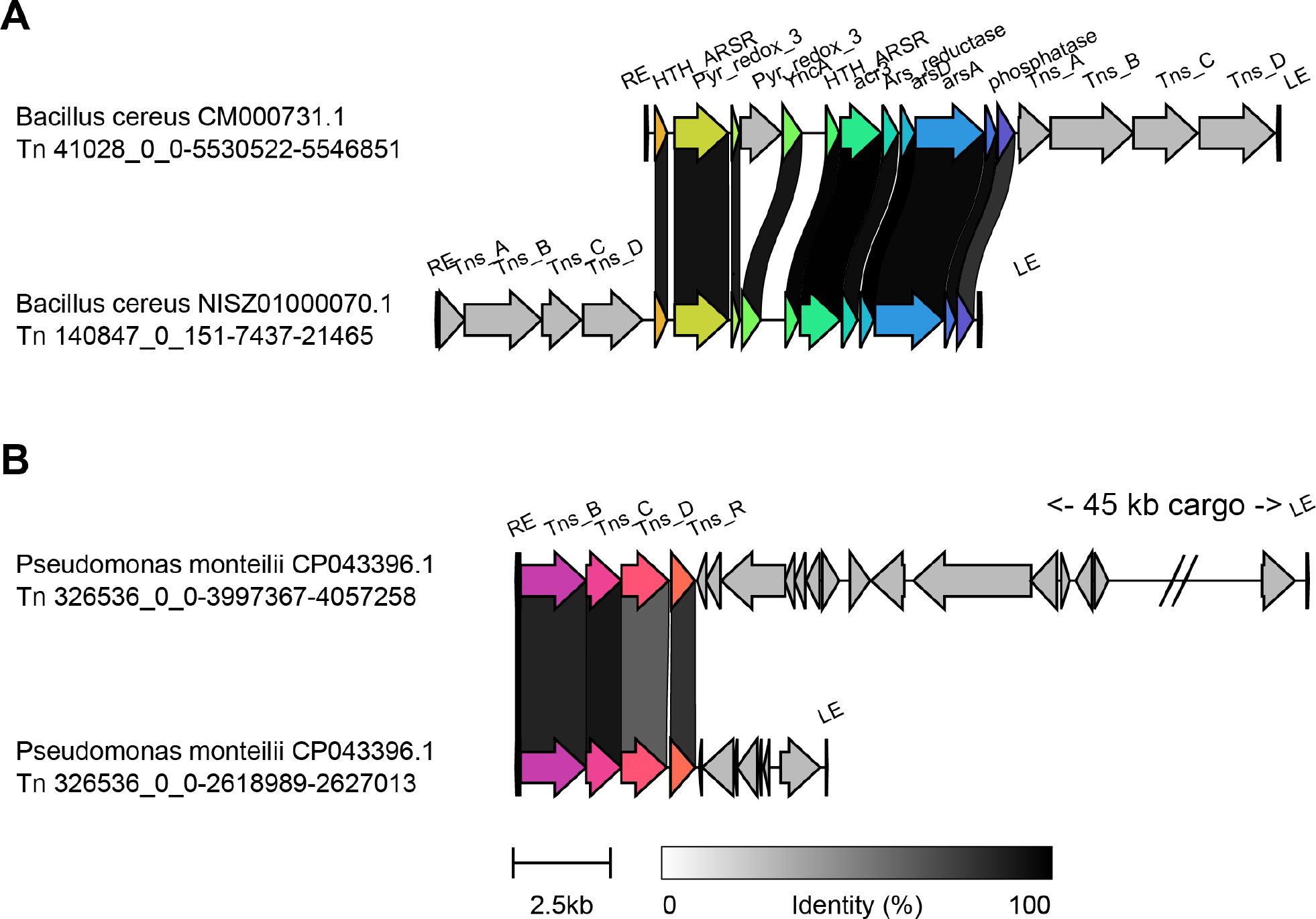
Closely related Tn7-like transposons possess different cargo, whereas same cargo can be mobilized by Tn7-like transposons from different clades. Transposons from clades 1 and 2 both mobilize a predicted arsenate resistance operon (**A**). The cargo encoded by two transposons from clade 3 integrated in the same genome mobilize different cargo repertoires (**B**).

**Figure S3.**
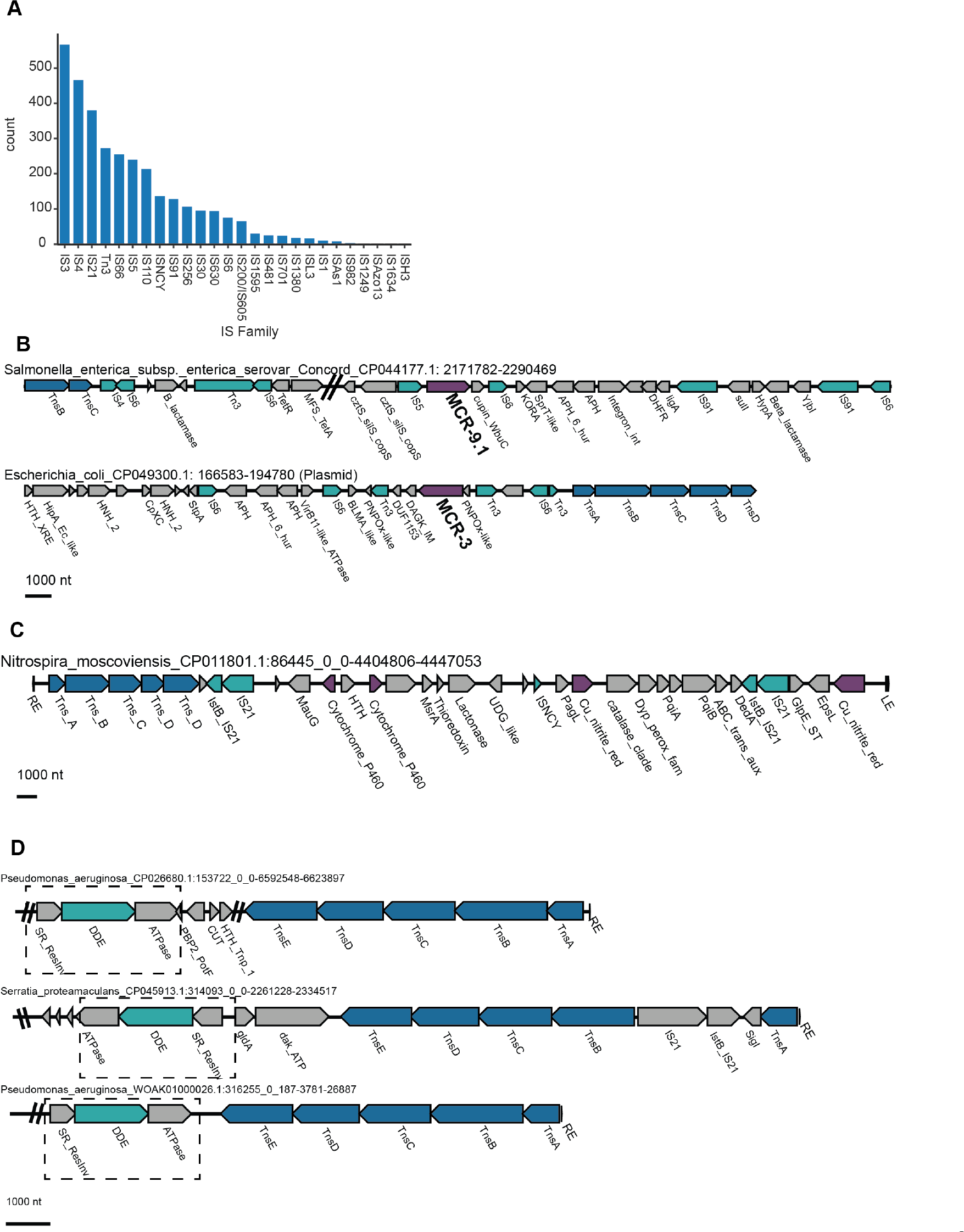
The cargo of Tn7-like transposons includes other MGEs. Histogram of transposons that harbor one or more insertion sequences (**A**) Examples of two colisitin- resistance genes that are part of a putative “nested” transposon (**B**). Two IS21-family transposons (teal) flank nitrogen-cycling genes (purple) (**C**). Unclassified, serine resolvase- carrying transposons are the cargo of multiple Tn7-like transposons (**D**).

**Figure S4.**
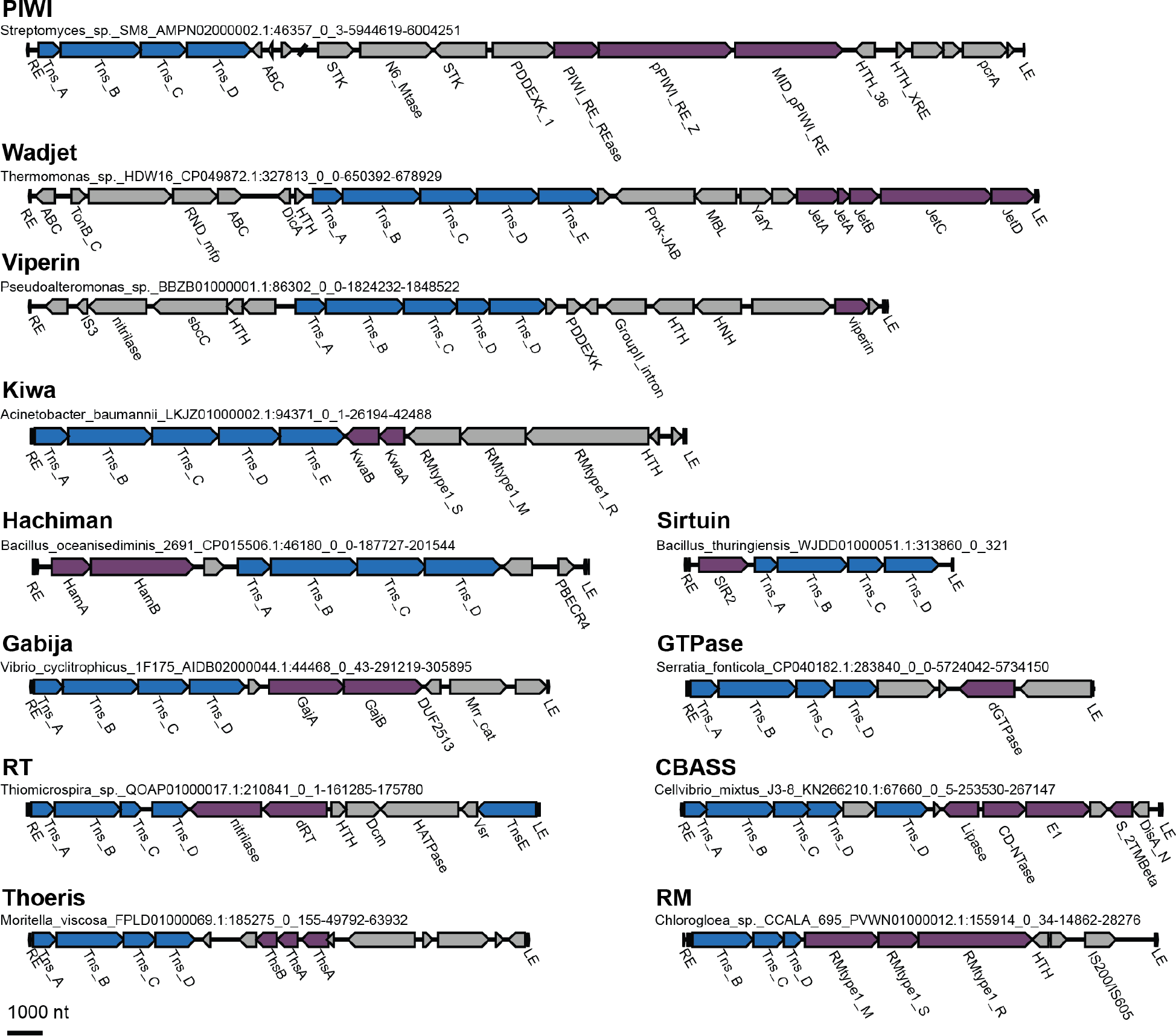
Innate immune systems mobilized by transposons. Selected examples of innate immune systems (purple) carried as cargo by Tn7-like transposons.

**Figure S5.**
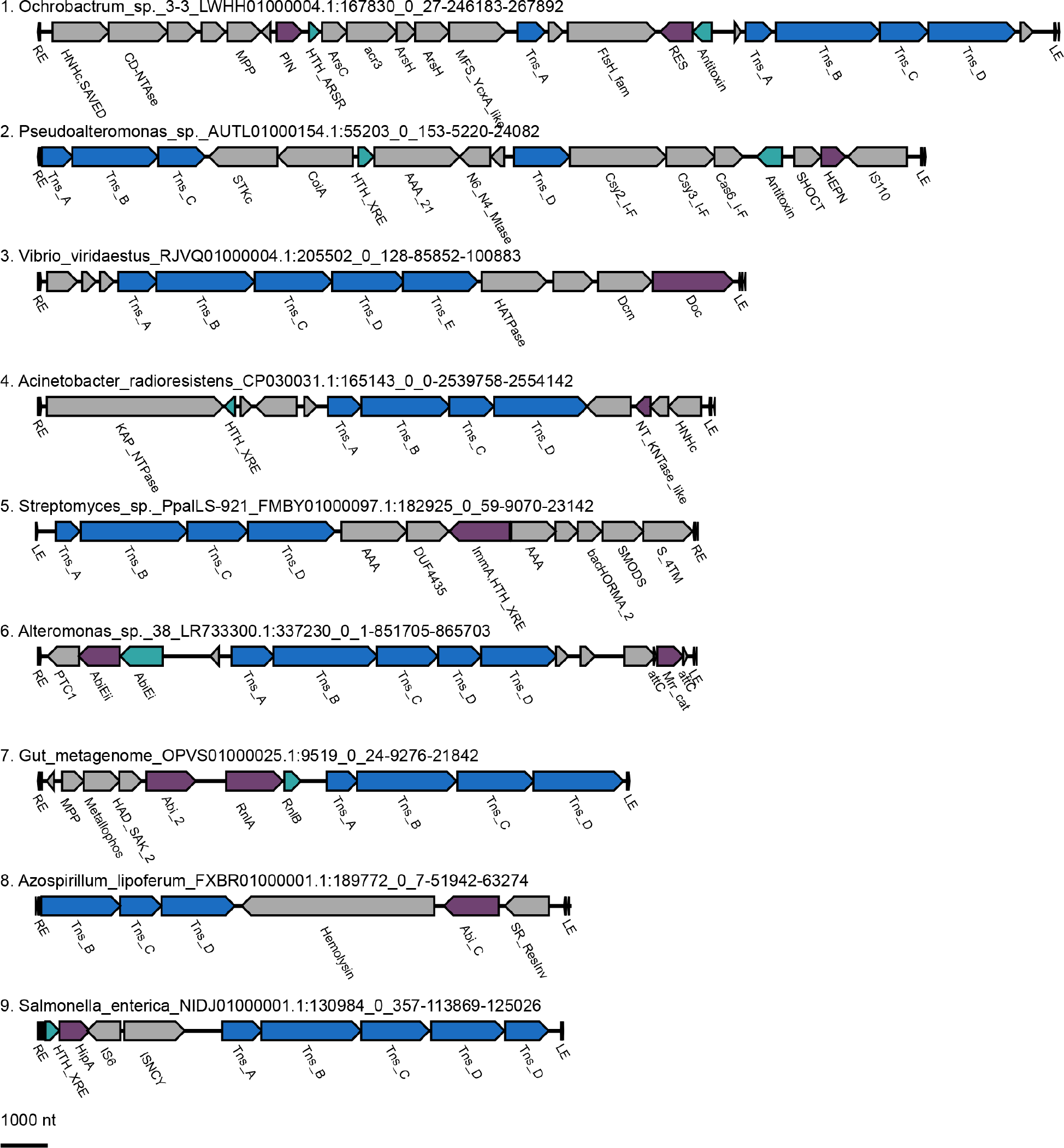
Toxin-antitoxin and abortive infection systems mobilized by transposons. Selected examples of toxins (purple) and antitoxins (teal) carried as cargo in Tn7-like transposons.

**Figure S6.**
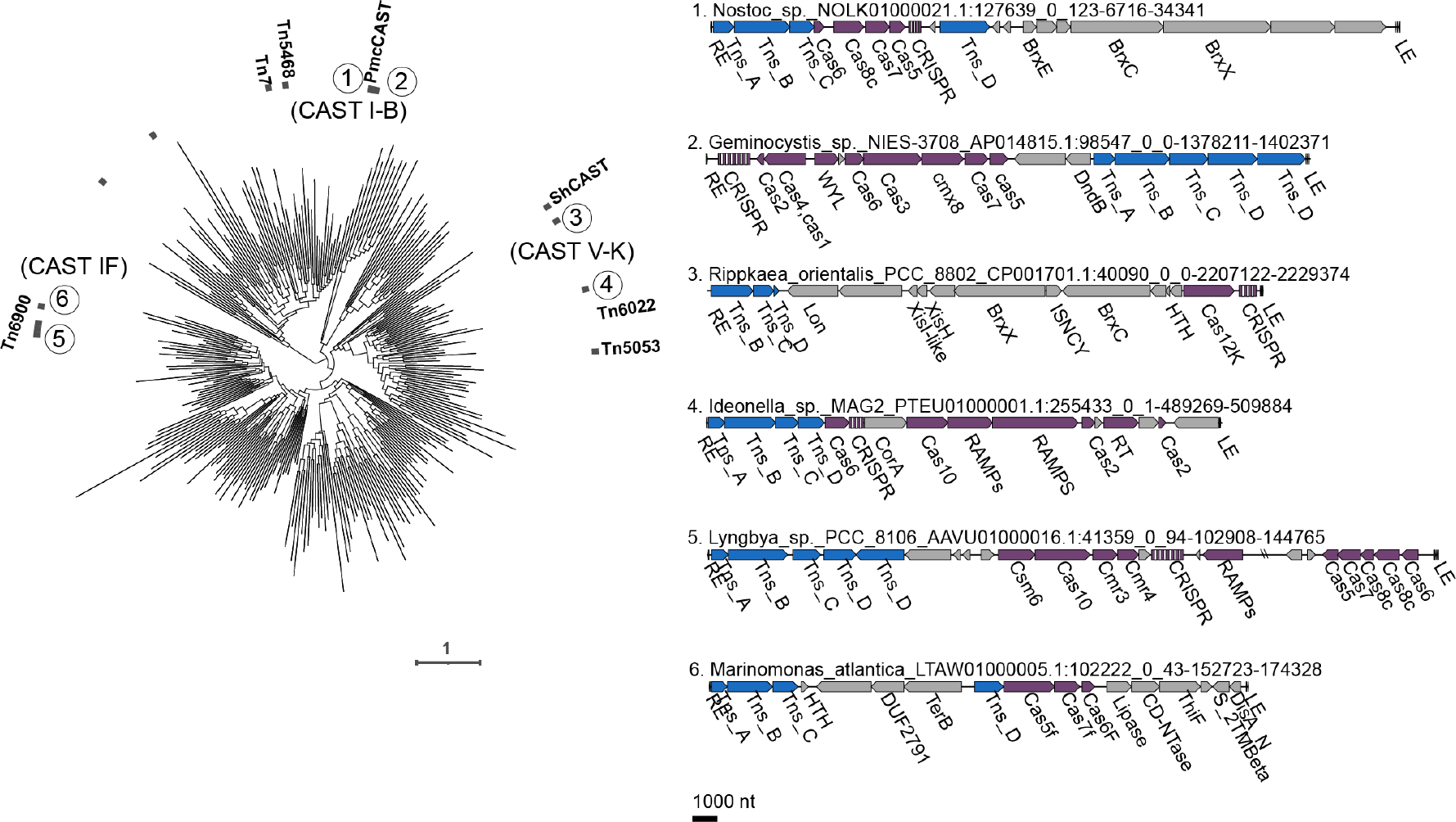
Capture of CRISPR-Cas systems by Tn7-like transposons. The presence of CRISPR arrays or *cas* genes are marked on the phylogenetic tree of TnsB. Six transposons are displayed, where the CRISPR-Cas components are highlighted in purple. Examples 1, 3 and 6 possess a CRISPR-Cas system functionally involved in transposition (see the main text). Examples 2, 4 and 5 carry an interference or adaptation module, whereas the most of the remaining transposons possess only standalone components of CRISPR-Cas (not shown).

**Figure S7.**
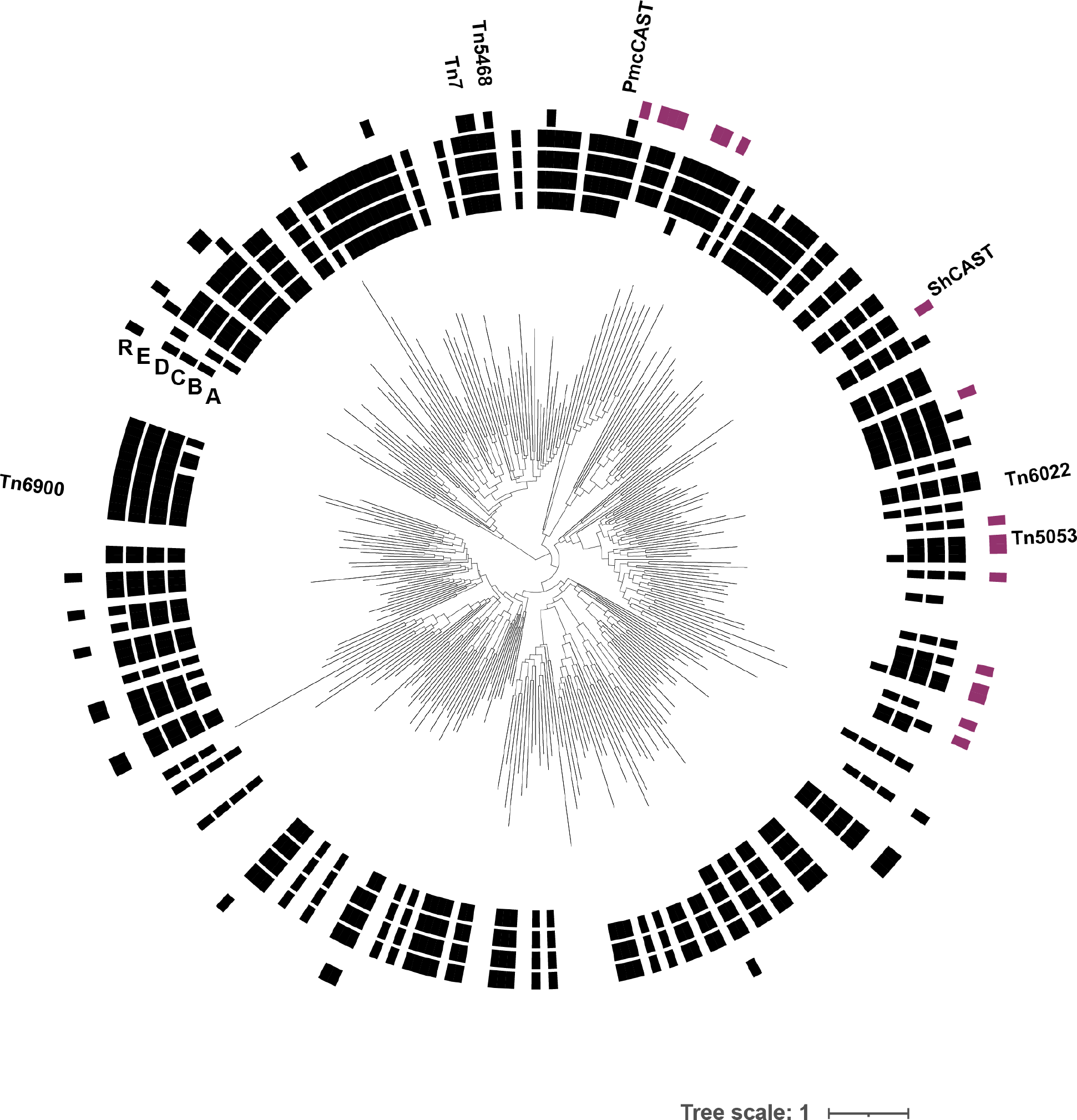
Replicative transposition is predicted by the absence of TnsA and presence of TnsR. The presence of core proteins in ≥50% of the transposons represented by each leaf are indicated on the outer rings of the TnsB tree. The resolvases of clades 2 and 3 are highlighted in purple, which do not cooccur with TnsA.

**Figure S8.**
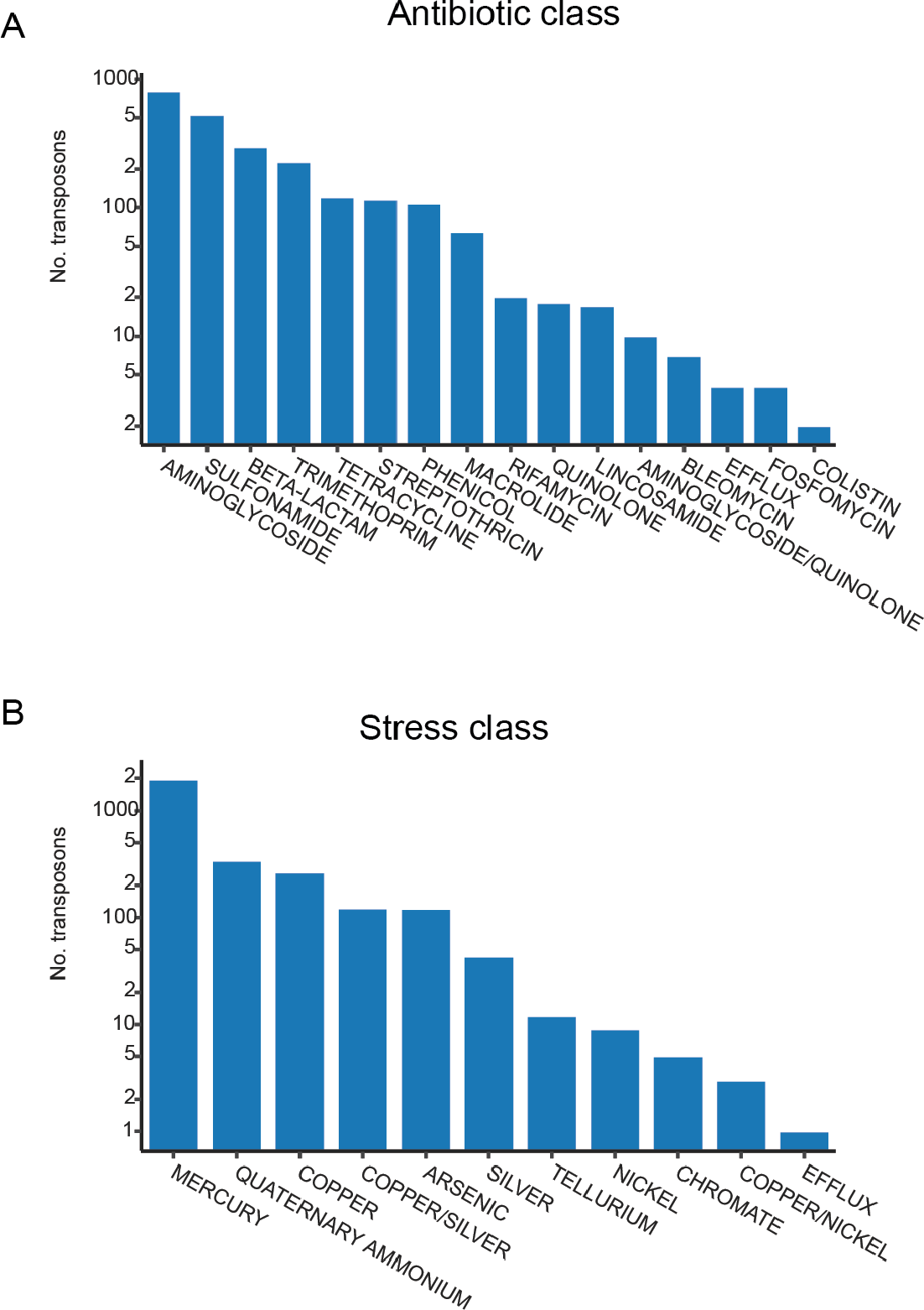
Antibiotic and stress resistance genes mobilized by Tn7-like transposons. Count of transposons harboring one or more antibiotic resistance genes (**A**) or stress-resistance genes (**B**). Note the log-scale.

**Figure S9.**
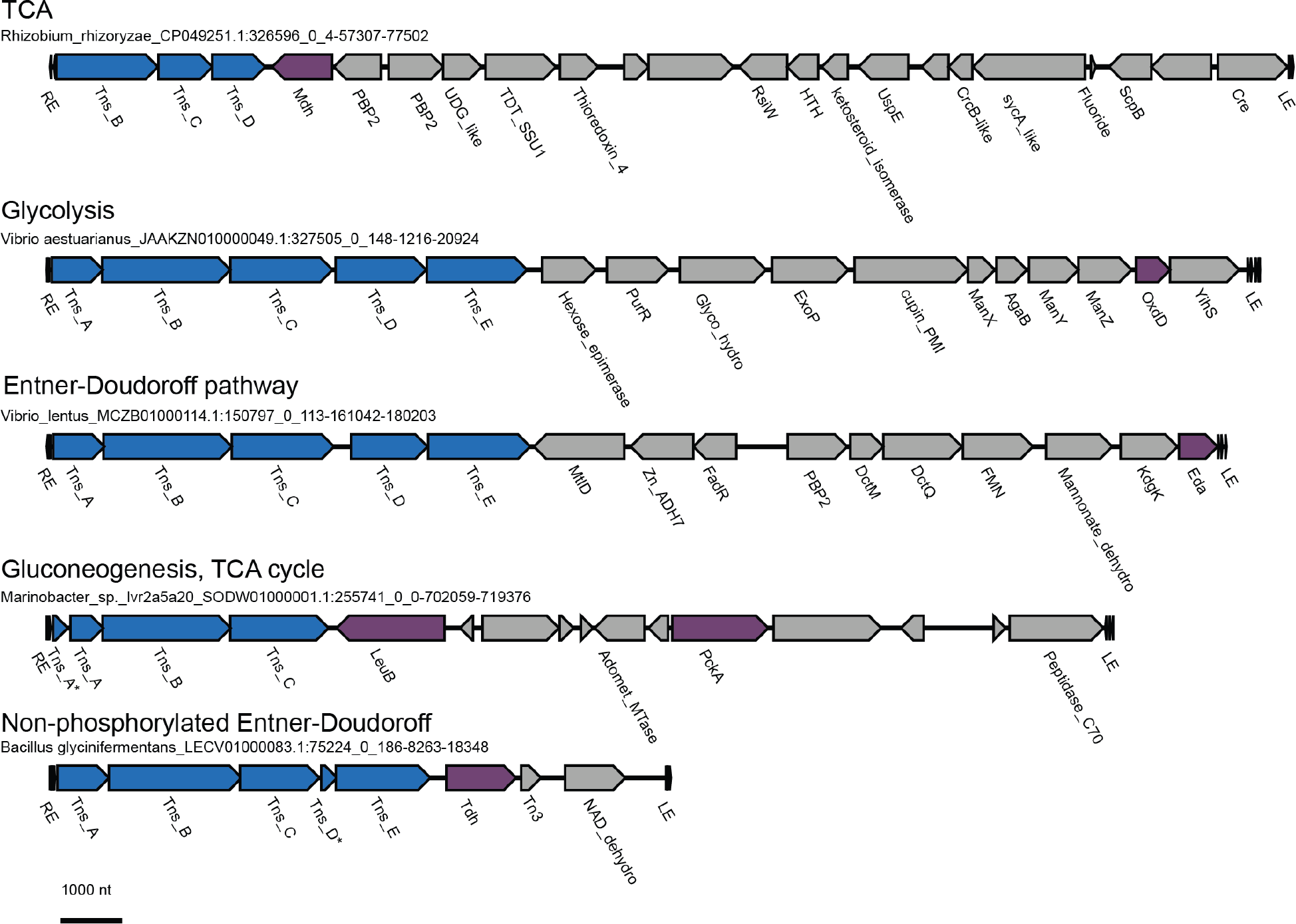
Central carbon metabolism genes mobilized by Tn7-like transposons. Central carbon metabolism genes are highlighted in purple, with their respective pathway labelled above each transposon. Abbreviations are as follows: Malate/lactate dehydrogenase (Mdh, COG0039), Oxalate decarboxylase (OxdD, COG2140), 2-keto-3-deoxy-6-phosphogluconate aldolase (Eda, COG0800), Isocitrate/isopropylmalate dehydrogenase (LeuB, COG0473), Phosphoenolpyruvate carboxykinase (PckA, COG1866), Threonine dehydrogenase (Tdh, COG1063).

## Supplement

**Table S1 – Metadata for all TnsB transposases.** For each TnsB protein, the host taxonomy and nucleotide coordinates are recorded. The leaf ID corresponds to the label of the newick tree. The “motif ID” column contains the Meme-formatted consensus sequence of the predicted TnsB binding site. Incomplete transposons whose ends could not be predicted do not have a motif.

**Table S2 – Metadata for all complete transposons.** The nucleotide coordinates and host DNA classification (chromosomal or plasmid) are reported for all complete Tn7-like transposons. Transposons were dereplicated (99% ID) and the representative sequence is provided for each transposon (“Tn_rep” column). The community to which each transposon belongs is listed for each hierarchical level.

